# Transcriptional Architecture of Synaptic Communication Delineates Cortical GABAergic Neuron Identity

**DOI:** 10.1101/180034

**Authors:** Anirban Paul, Megan Crow, Ricardo Raudales, Jesse Gillis, Z. Josh Huang

## Abstract

Understanding the organizational logic of neural circuits requires deciphering the biological basis of neuron type diversity and identity, but there is no consensus on defining a neuron type. We analyzed single cell transcriptomes of anatomically and physiologically characterized cortical ground truth populations and conducted a computational genomic screen for transcription profiles that distinguish them. We discovered that cardinal GABAergic neuron types are delineated by a transcriptional architecture that encodes their synaptic communication patterns. This architecture comprises 6 categories of ~40 gene families including cell adhesion molecules, transmitter-modulator receptors, ion channels, signaling proteins, neuropeptides and vesicular release components, and transcription factors. Combinatorial expression of select members across families shapes a multi-layered molecular scaffold along cell membrane that may customize synaptic connectivity patterns and input-output signaling properties. This molecular genetic framework of neuronal identity integrates cell phenotypes along multiple axes and provides a foundation for discovering and classifying neuron types.

## INTRODUCTION

Since the discovery that individual neurons are basic building blocks of the nervous system (Cajal, 1892), the immense diversity and heterogeneity of nerve cells have remained a formidable challenge for deciphering the organizational logic of neural circuits (Armananzas and Ascoli, 2015; Bota and Swanson, 2007). Recent technical advances have accelerated progress in anatomical, physiological, developmental and functional studies that increasingly reveal multilayered and multi-dimensional variations of neuronal phenotypes and properties (Huang and Zeng, 2013; Luo et al., 2008). A fundamental question of broad significance is whether these variations are continuous, largely subjective to measurements, and can only be managed by empirical and operational grouping, or whether multiple distinct and congruent cell features can be integrated to define and classify discrete “cells types” that reflect biological reality and mechanisms (DeFelipe et al., 2013; Seung and Sumbul, 2014). The problem of neuronal diversity and census are unlikely to be solved without solving the equally if not more fundamental problem of neuronal identity, the flip side of the cell type coin (Seung and Sumbul, 2014). However, in many brain regions, such as the cerebral cortex, there is no consensus on what a neuron type is; the biological basis of neuronal identity is poorly understood and the classification scheme of cell types remain contentious (Battaglia et al., 2013; DeFelipe et al., 2013; Petilla Interneuron Nomenclature et al., 2008).

As individual neurons also constitute basic units of gene regulation in the brain, a major determinant of each neuron’s characteristic phenotype and function likely lies in its transcription program, shaped by its chromatin landscape customized from the genome. Recent advances enable mRNA sequencing of individual cells (Tang et al., 2009), and several studies have aimed to discover and classify neuron types using high-throughput single cell RNAseq (scRNAseq) and statistical clustering (Macosko et al., 2015; Tasic et al., 2016; Usoskin et al., 2015; Zeisel et al., 2015). A major challenge has been to map transcriptome-based statistical cell clusters, which are prone to technical noise and methodological bias, to the biological ground truth of cell types - their anatomical and physiological properties that constrain and contribute to their function in neural circuits. In the retina, where cell types are among the best understood in the mammalian nervous system, high throughput scRNAseq has identified transcriptionally distinct cell population markers that correlate to known types and suggested novel candidate types (Macosko et al., 2015; Shekhar et al., 2016). In the cerebral cortex, where cell type definition is often ambiguous and controversial, scRNAseq analyses have parsed cells into multiple “transcriptional types” (Tasic et al., 2016; Zeisel et al., 2015), but the boundaries of such statistical types often appear fluidic if not problematic, and the extent to which they correlate to bona-fide biological types jointly defined by anatomical and physiological features remain unclear. Thus although scRNAseq allows comprehensive, quantitative and high throughput measurements of gene expression, a fundamental unresolved issue is whether and how transcription profiles might contribute to the molecular genetic root of neuron types. Discovering such transcriptional basis of neuronal identity is prerequisite for using a transcriptome-based approach to decipher neuronal diversity and enumerate cell census.

Beyond cell type discovery and classification, a major promise of transcriptome analysis is to uncover the molecular mechanisms that underlie multi-faceted yet functionally congruent cell phenotypes and properties. Although an increasing set of molecular markers have been identified for different cell populations (Shekhar et al., 2016; Tasic et al., 2016; Zeisel et al., 2015), comprehensive and high-resolution molecular portraits that mechanistically and coherently explain and predict cell phenotypes have yet to be achieved.

Here, we have discovered the transcription architecture underlying the core identity of cardinal GABAergic neuron types in the cerebral cortex. Unlike several recent studies that classify neurons using unsupervised statistical clustering of single cell transcriptomes from unbiased populations (Zeisel et al., 2015) or relatively broad populations (Tasic et al., 2016), we analyzed the transcriptomes of ~530 GABAergic neurons in mature mouse neocortex derived from 6 cardinal types or subpopulations and that were captured by intersectional or lineage-based genetic labeling. Using these anatomy and physiology defined ground truth populations as an assay, we designed a supervised and machine learning-based computational genomics strategy to screen through each of the ~620 HGNC (Human Genome Nomenclature Committee) annotated gene families for those whose differential expression among family members reliably distinguish these subpopulations. Remarkably, approximately 40 gene families implicated in regulating synaptic connectivity and communication best distinguish these subpopulations. These gene families constitute 6 functional categories that include cell adhesion molecules, neurotransmitter and modulator receptors, ion channels, membrane-proximal signaling molecules, neuropeptides and vesicular release components, and transcription factors. Combinatorial and coordinated expression of select family members across functional categories shapes a multi-layered molecular scaffold along the cell membrane that appears to customize the pattern and property of synaptic communication for each cell population. We further provide evidence that expression profiles of transcription factors register the developmental history of GABAergic neurons and contribute to the concerted gene expression patterns that shape cell phenotypes. These findings suggest that neuron type identity is encoded in a transcriptional architecture that orchestrates functionally congruent expression across multiple gene families to diversify and customize the patterns and properties of synaptic communication. This overarching and mechanistic definition of neuron type integrates, explains and predicts cell phenotypes along multiple axes and provides an intellectual framework for neuron type discovery and classification in the nervous system.

## RESULTS

### Single cell transcriptomes of ground truth GABAergic cell types and subpopulations

Our overall strategy in exploring the molecular basis underlying cortical GABAergic neuron identity is to examine and compare high resolution transcription profiles of a set of well characterized cell types or subpopulations defined by multiple anatomical, physiological and developmental attributes (He et al., 2016; Taniguchi et al., 2011). Cortical GABAergic neurons can be parsed into several broad classes, non-overlapping populations and, in a few cases, bona-fide types based on developmental origin, innervation targets, and molecular markers (Kepecs and Fishell, 2014; Somogyi et al., 2014). The embryonic medial and caudal ganglionic eminences (MGE and CGE) give rise to two broad groups, the former is divided into parvalbumin (PV) and somatostatin (SST) populations and the latter is marked by 5HTR3a (Rudy et al., 2011) (Figure 1A-B). The PV population includes fast-spiking basket cells (PVBC) that innervate the perisomatic region (Hu et al., 2014) and chandelier cells (ChC) that target the axon initial segment (AIS) (Somogyi, 1977; Taniguchi et al., 2013). The SST population includes Martinotti cells (MNC) that target distal dendrites (Wang et al., 2004), long projection cells (LPC) (Tamamaki and Tomioka, 2010) and multiple other cell types. The 5HTR3a group includes the Vassoactive intestinal peptide (VIP) and Reelin populations, and the VIP population comprises interneuron-selective dis-inhibitory cells (ISC) (Pi et al., 2013; Staiger et al., 2004), Cholecystokinin (CCK) small basket cells (CCKC) (Armstrong and Soltesz, 2012; Freund and Katona, 2007) and likely additional cell types. Accumulated anatomical, physiological, and molecular evidence indicate that these are non-overlapping subpopulations, and ChC, LPC and PVC are considered cardinal types (He et al., 2016).

**Figure 1.**
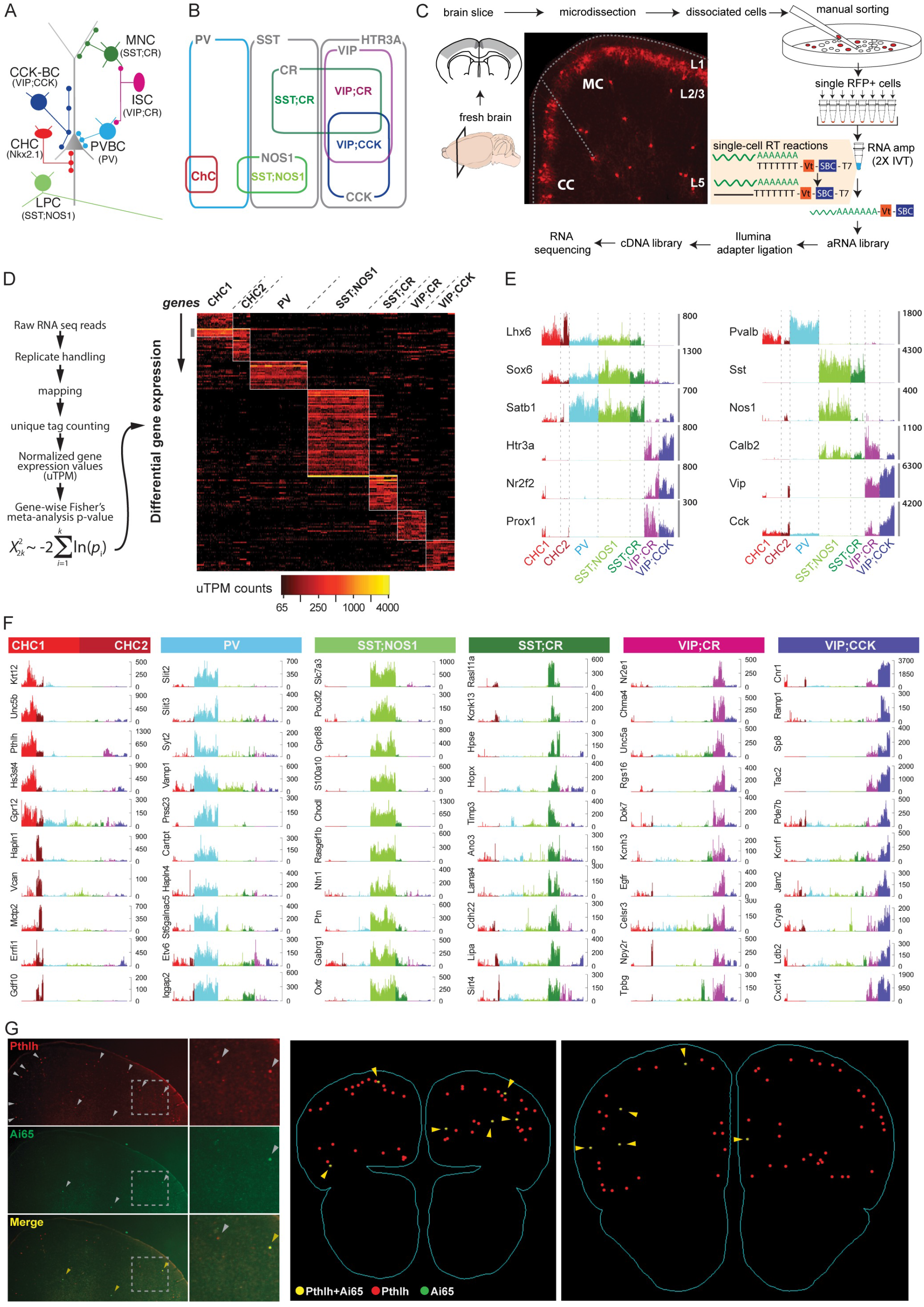
Workflow of transcriptomic analysis of cortical GABAergic ground truth populations. **(A)** Schematic of 6 major neocortical GABAergic cell types with characteristic cellular and subcellular innervation patterns. ChC: chandelier cells, PVC: PV basket cells, LPC: long projection cells, MNC: Martinotti cells, ISC: interneuron selective cells, CCKC: CCK basket cells. Molecular markers that label or include these cell types are shown in parenthesis. Combinatorial driver lines that capture each population are shown in Figure S1. **(B)** Cortical GABAergic neurons comprise 3 non-overlapping populations labeled by PV, SST and 5HTR3A. Within these, 6 ground truth populations (GTPs) are labeled by lineage or combinatorial markers that include bona-fide cell types shown in (A). **(C)** Experimental workflow. From transgenic mice containing genetically labeled GTPs, fresh coronal brain sections containing cingulate, somatosensory and motor areas were microdissected and dissociated to single cell suspension. Individual RFP-labeled neurons were manually sorted. Single neurons were reverse transcribed with unique molecular identifiers and pooled for amplification using two rounds of in-vitro transcription followed by multiplexed cDNA library generation and Illumina sequencing. **(D)** Bioinformatics pipeline showing read mapping, replicate handling, duplicate read elimination by counting unique tags, normalization and deriving Fisher’s meta-analytic p-value for differentially expressed (DE) transcripts in each GTP. DE heatmap shows 183 genes across GTPs as absolute reads or, unique transcripts per million (uTPM) counts. ChCs are split into layer 2/3 (CHC1) and layer 5/6 (CHC2) groups. **(E)** Barplots showing uTPM values of known markers for individual cells from each GTP (colored): Lhx6, Sox6 and Satb1, for MGE and Htr3a, Nr2f2, Prox1, for CGE populations (left). Expression of other markers used for combinatorial targeting of GTPs matched perfectly in expected cell populations (right). Y-axis denotes unique mRNA copies in uTPM. **(F)** Barplot of novel cell type markers for each GTPs; y-axis: uTPM counts. Note that CHC1 and CHC2 express distinct molecular markers. **(G)** Fluorescent mRNA in-situ hybridization showing that a candidate CHC1 marker Pthlh (grey arrowheads) co-localizes with genetically labeled CHCs (tdTomato reporter gene in a *Nkx2.1;Ai14* mouse tamoxifen-induced at E17.5, yellow arrowheads) (left). Two representative sections from serial 3D reconstruction of the forebrain of show >95% Ai14 cells are Pthlh positive (right). Additional in-situ data in Figure S1.

We have developed combinatorial Cre and Flp recombinase driver lines to capture 6 GABAergic subpopulations and cell type through the activation of Ai14 or Ai65 reporters that express the fluorescent protein tdtomato (RFP) (He et al., 2016; Taniguchi et al., 2011): 1) The *Nkx2.1-CreER* driver allows lineage and birth timing based targeting of ChCs, 2) the *PV-Cre* driver labels a broad class of fast-spiking basket cells, 3) the *SST-Flp;nNOS-CreER* drivers target a highly unique type of long-projecting GABAergic neurons, 4) the *SST-Flp;CR-Cre* drivers include Martinotti cells and likely other cell types, 5) the *VIP-Flp;CR-Cre* drivers include interneuron-selective cells and likely other cell types, 6) the *VIP-Flp;CCK-Cre* drivers include CCK basket cells and likely other cell types. Together, we define these 6 populations as Ground Truth Populations, or GTPs.

Using manual sorting (Paul et al., 2012; Sugino et al., 2006) of single RFP-labeled cells from microdissected motor and somatosensory cortical slices from mature (6 weeks old) mice (Figure 1C), we obtained high depth transcriptome of ~584 cells from the 6 GTPs (Figure S1D; see Materials and Methods). This unique dataset thus contains high-resolution transcriptomes of phenotype-defined cortical GABAergic GTPs. Compared with previous 6bp UMI-based method (1.8-4.7K genes, (Zeisel et al., 2015) and non-UMI RPKM based readouts (7.2K genes, (Tasic et al., 2016), our method of manual sorting coupled with linear amplification (Eberwine et. al. 1992) with 10bp UMIs improved single cell gene detection and quantification (~10K genes; Figure S1H). Compared with DropSeq which allows vast throughput at low cost (Macosko et al., 2015), our complementary approach achieves more comprehensive and quantitative transcriptome measurement of targeted cell populations, which facilitates more in-depth analysis of molecular profiles that may contribute to cell phenotypes and identity.

Differential expression (DE) analysis revealed 190 genes that were differentially expressed among GTPs with each single cell expressing >50uTPM, >4 folds enrichment and with p-value < 5X10^-4^ (Figure 1D and Table S1). We detected between 26-91 DE genes for each GTP population. A subset of these DE genes is shown in Figure 1F as single cell barplots. We confirmed the expression of multiple known markers for MGE (Lhx6, Sox6, and Satb1) and CGE (Htr3a, Nr2f2, and Prox1) derived interneurons and all markers used for combinatorial targeting matched perfectly to appropriate cell populations (Figure 1E), validating our method and dataset. To explore the laminar distinction of GTPs, we profiled *Nkx2.1-CreER* labeled ChCs from upper (L1-L2 boundary, CHC1) and deeper (L5+6, CHC2) layer cohorts. Although CHC1 and CHC2 transcriptomes were highly similar, we detected ~11 genes that were enriched in CHC2 (Figure 1 D and F). We validated the GTP specific expression of ~10 selected transcripts using fluorescent double mRNA in-situ hybridization in appropriate driver lines in which a GTP can be detected with a RFP mRNA in situ probe (Figure 1G and Figure S2A). In particular, we discovered a putative pan-CHC transcript Pthlh: ~95% of Ai14-labeled CHCs were positive for Pthlh (136/143 cells) and their laminar distribution recapitulate ChC pattern in adult frontal cortex (Taniguchi et al., 2013) (Figure 1G, Figure S2A).

Previous DE analyses often reveal molecular markers that, although useful, appear piece meal and do not readily inform or explain cell properties (Tasic et al., 2016; Zeisel et al., 2015). To systematically examine the relationship between differential gene expression and cell phenotypes, we analyzed whether and how functional gene ensembles (e.g. gene families) relate to cellular properties of GTPs.

### A computation genomic screen identifies gene families and categories that distinguish and characterize GTPs

Cellular properties (e.g. fast spiking) emerge from operations of macromolecular machineries (e.g. sodium and potassium channel complexes consisting of multiple interacting core subunits, auxiliary subunits, scaffolding proteins); each component is often implemented as one of multiple variants encoded by a gene family (e.g. I of 9 members in the Nav family). Thus variations of cell properties (e.g. spike width) among cell types often result from differential usage or expression levels of select members (e.g. Nav.1 vs Nav1.6) with characteristic biochemical and biophysical properties that confer customized properties to modular cellular machines (Hartwell et al., 1999). Given the highly distinct and well-characterized anatomical and physiological features among GTPs, we hypothesized that these phenotypic differences result from systematic and coherent transcriptional differences across multiple gene families of different functional categories, much beyond a piece meal set of serendipitous markers. The unique strength of our experimental design, whereby single cell transcriptomes derive from 6 GTPs, provided a powerful *assay* to systematically *screen* for such functional gene ensembles that distinguish and characterize GTPs. To efficiently and comprehensively identify such gene families, we designed a supervised, machine learning based algorithm, MetaNeighbour (Crow et. al. 2017), to screen all the Gene Ontology (GO) terms and all the ~620 annotated gene families.

The essence of our computational genomics screen is to detect whether a given set of genes (e.g. gene families) shows preferentially correlated expression among cells known to possess the same identity (Figure 2A). Because our single cell transcriptomes derive from 6 GTPs, this data structure allowed us to characterize the similarity between all pairs of single cells using covariation of expression level in many known gene sets and measure whether a given gene set correctly links cells of known identity. In a network formalism, each cell is a node and cells are linked as probabilistically related based on the similarity (correlation) of their transcriptional profiles across a given set of genes (Figure 2A). This network can be used to classify cells based on their proximity within it: cells which are close within the network are predicted to share an identity (see Methods). A subset of the GTP labels are applied to cells, giving a sub-network of cells with known identities which can classify unlabeled cells. We then hold back the GTP identity of some cells (cross-validation) and attempt to predict their identities using this subnetwork of known identities. A cell is predicted to have a given identity if its neighboring cells (grouped by similarity in their gene set expression) belong to a sub-network that defines that identity (Figure 2A). We report on the efficacy of this test using mean area under the receiver operator characteristic curve (AUROC), which maps to the probability that the assignment is correct, if it was making a single binary (positive/negative) choice (Figure 2A). Having constructed a computational assay for cell identity, we vary the transcriptomic features (e.g. gene families) used to characterize cells as neighbors of one another. This computation screen thus selects functional gene ensemble features (e.g. gene families) which jointly distinguish cell identities. We perform a stratified cross-validation which allows us to explicitly block technical sources of variation in single-cell analysis (see Methods), in close parallel to our meta-analytic evaluation of single-cell data (Crow, 2016).

**Figure 2.**
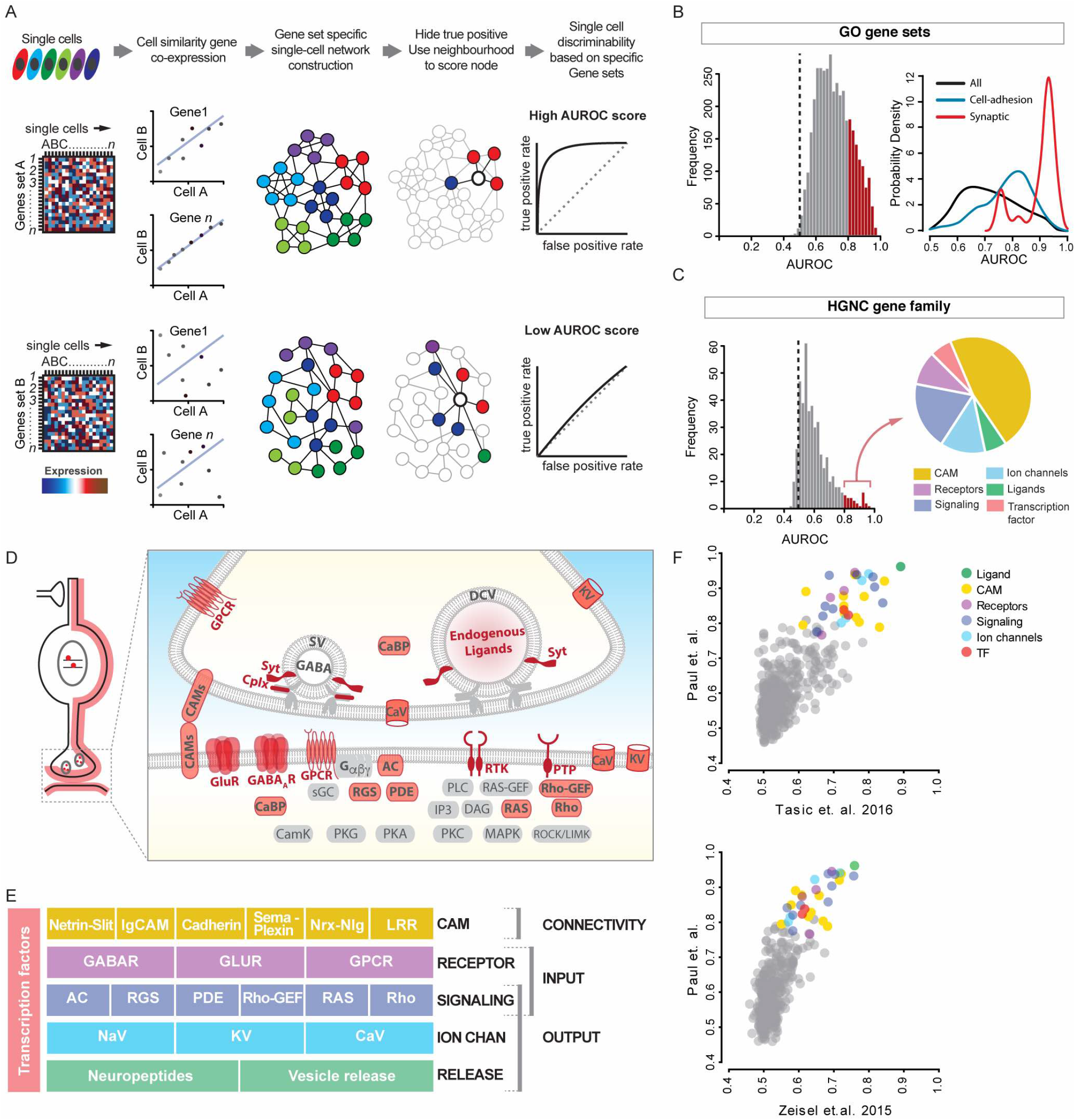
A computation genomic screen identifies gene families and categories that distinguish and characterize GTPs. **(A)** Schematic of bioinformatic pipeline for MetaNeighbor. Single cells gene expression values for gene ensembles (e.g. gene sets A and B) are used to construct cell-cell networks based on similarity of co-expression, such that cells similar in gene expression space are close neighbors shown by connecting lines. Cell identity (i.e. GTP identity) labels inherent to single cells (shown as colors) are then withheld and its identity inferred based on connectivity to its immediate neighbors. The aggregate probability of being identified as the correct GTP is reported as AUROC (area under receiver operator curve) score - the probability that a cell was correctly assigned to its GTP identity, where AUROC=0.5 represents chance performance. Depending on the gene set used some networks perform better in the classification task. By iterating the process through collection of gene sets grouped by gene ontology (GO) or Human Genome Nomenclature Consortium (HGNC) families or custom categories, a ranked list of AUROC for all such gene sets is generated. **(B)** Left: Frequency distribution histogram of AUROC values of ~3800 GO terms. Red bars indicate GO-terms that performed with AUROC>0.8. Black dotted line indicate chance performance (AUROC=0.5). Right: a probability density function plot showing GO-terms containing keyword “synaptic”(red) and “cell-adhesion”(green) have rightward skewed distributions of AUROC>0.8 compared to the probability density of all terms (black). **(C)** Left: Histogram of 442 HGNC gene families shows ~7% have a GTP identity prediction value exceeding AUROC=0.8. Right: High-performance gene families with AUROC>0.8 mainly comprise six gene categories. **(D)** Schematic showing that gene products encoded by the high-performance gene families (except transcription factors) primarily localize along cell and synaptic membrane; they mainly contribute to synaptic connectivity and signaling. **(E)** High-performance gene families constitute 5 layers of functional categories that organize synaptic connectivity and input-output signaling. **(F)** MetaNeighbor analysis of two independent single cell transcriptome datasets yields the same rank order of HGNC gene families, suggesting that these gene sets are a fundamental feature of GABAergic neuron identity. Abbreviations: GPCR, G-protein coupled receptor; SV, synaptic vesicle; DCV, dense core vesicle; KV, voltage-gated potassium channel; CaV, voltage-gated Ca+2 channel; Syt, Synaptotagmins; Cplx, Complexins; CaBP, Calcium binding protein; CAMs, cell adhesion molecules; GluR, glutamate receptor; GABA_A_R, GABA-A receptor; Gα,β,γ, Trimeric G-protein subunits α,β,γ; AC, Adenylate cyclase; PTP, Protein tyrosine phosphatase, sGC, soluble Guanylate cyclase; RGS, Regulator of G-protein signaling; PDE, Phosphodiesterase; PLC, Phospholipase C, RAS-GEF, Ras Guanine nucleotide exchange factor; Rho-GEF, Rho Guanine nucleotide exchange factor; IP3, Inositol triphosphate; DAG, Diacyl glycerol; RAS, Ras GTPase; Rho, Rho GTPase; CamK, Calcium/Calmodulin dependent protein kinase; PKG, Protein kinase G, PKA, Protein kinase A, MAPK, Mitogen activated protein kinase, ROCK/LIMK, Rho associated kinase/ LIM kinase.

We first screened for gene ensembles according to GO terms, using both randomized labels (AUROC~0.5) and randomized gene sets as controls. Among the GO terms, those containing the keyword “synaptic” gave the highest AUROC score ranging between 0.91-0.98, suggesting that genes implicated in synaptic connectivity and function are most discriminating for GTPs (Figure 2B, Table S3). Although informative, GO terms are too broad and redundant for describing neuronal phenotypes and properties. To identify more specific and extensive gene categories, we screened through all gene families annotated in the Human Genome Nomenclature (HGNC) database (see Methods). We identified ~40 gene families (i.e. 7% of all gene families) with AUROC scores >0.75, generally regarded as a stringent threshold (Figure 2C, Table S4). Strikingly, these gene families all fell into only 6 functional categories (Figure 2C-D): 1) cell adhesion molecules, 2) receptors for neurotransmitters and modulators, 3) voltage-gated ion channels, 4) regulatory signaling proteins, 5) neuropeptides and vesicle release machinery, 6) transcription factors. It is immediately evident from this list that except transcription factors (TFs), all other gene categories encode proteins that localize along or close to cell and synaptic membrane (Figure 2D) and contribute to a singular aspect of neuronal biology - synaptic communication, which is implemented through synaptic connectivity and input-output signaling properties (Figure 2E).

To validate this discovery, we applied the MetaNeighbour screen to two independent scRNAseq datasets from equivalent cell populations (Tasic et al., 2016; Zeisel et al., 2015). Despite notable differences in experimental design, RNA amplification, library construction and mapped read tallying, our meta-analysis (Crow, 2016) of the combined dataset from the three studies validated all of our ground truth populations and 46% of the published data (18/39 transcriptional types: 7/16 from Zeisel et al., 2015 and 11/23 from Tasic et al., 2016) (Figure 2F). More importantly, we found nearly identical results on the rank order of gene families that best discriminate equivalent cell populations (i.e. 6 GTPs) in the three dataset, and the AUROC values of all the GTP-distinguishing gene families (Figure 2C) were well correlated in pair wise comparisons even though the scores from the other two datasets were modestly lower (Figure 2F).

Together, our results indicate that, among the ~20,000 protein-coding genes constituting ~620 gene families in mouse genome (442 HGNC families with 3 or more members were analyzed), GABAergic GTPs can be effectively distinguished by a small fraction of ~40 families constituting 6 functional categories. These gene categories appear to construct a coherent transcriptional architecture encoding a 5-layered molecular scaffold along the cell membrane that organizes and customizes synaptic connectivity and input-output signaling. This result thus suggests that the core identity of GABAergic neurons might be encrypted in key transcription features that coordinate two fundamental cell attributes - the pattern and style of synaptic communication.

In the following sections, we examine each of the 6 gene categories and demonstrate how coordinated expression of select members across families and categories correlate with, contribute to and predict cell phenotypes and properties that together shape the identities of GTPs.

### Differential expression of cell adhesion molecules and carbohydrate modifying enzymes among GTPs suggests large capacity for cell surface and extracellular matrix labels

Each GABAergic neuron receives hundreds to thousands of inputs from diverse presynaptic neurons and in turn contacts similar number of postsynaptic neurons of multiple types (Figure 2A). These synaptic connections are established between specific cell types and at designated subcellular locations (i.e. wiring specificity) (Huang et al., 2007) and are further customized in their transmission properties for specific pre- and post-synaptic partners (i.e. synapse specificity) (de Wit and Ghosh, 2016). Classic studies have postulated a large set of “individual identification tags” on cell surface that allow neurons to distinguish one another and selectively connect to appropriate partners (Sperry, 1945). Studies in past decades have identified dozens of gene families encoding hundreds of neuronal cell adhesion molecules (CAMs) and synaptic adhesion molecules, some with thousands of splice variants, suggesting a molecular basis for the capacity and diversity of cell surface tags (Figure 2B) (de Wit and Ghosh, 2016; Kolodkin and Tessier-Lavigne, 2011). Through combinatorial ligand-receptor signaling, these CAMs play specific and overlapping roles during neural circuit assembly, including axon guidance, neurite branching and pruning, cellular and subcellular recognition, synapse formation and specificity, synapse property and plasticity. It is unclear to what extent these same adhesion molecules are reused in mature neurons to maintain cell morphology and connectivity, and to regulate synaptic transmission and plasticity. In particular, the repertoires of CAMs expressed in specific cell types in mature circuits are unknown (de Wit and Ghosh, 2016).

Our computation genomics screens identified multiple CAM gene families that effectively discriminate GTPs (Figure 2D-E). Based on these broadly annotated HGNC families (total of ~660 genes) and neurobiology literature (de Wit and Ghosh, 2016; Kolodkin and Tessier-Lavigne, 2011; Takahashi and Craig, 2013), we selected a set of ~275 genes encoding all major neuronal CAMs and organized them into 12 adhesion groups according to sequence homology and receptor-ligand relationships (Figure 3B; Table S5; See Methods and Supplemental Text). Notably, nearly all major groups of neuronal CAMs implicated in different aspects of neuronal development are expressed in GTPs, and each GTP on average expresses ~200 genes encoding CAMs (Figure 3C). This was an underestimate of CAM diversity as our RNAseq method does not detect splicing variants. Among the total ~275 neuronal CAM genes, 130 show highly distinct subpopulation profiles (Figure 3E, TableS2). Strikingly, multiple CAM families each manifests differential expression among GTPs (Figure 3F).

**Figure 3:**
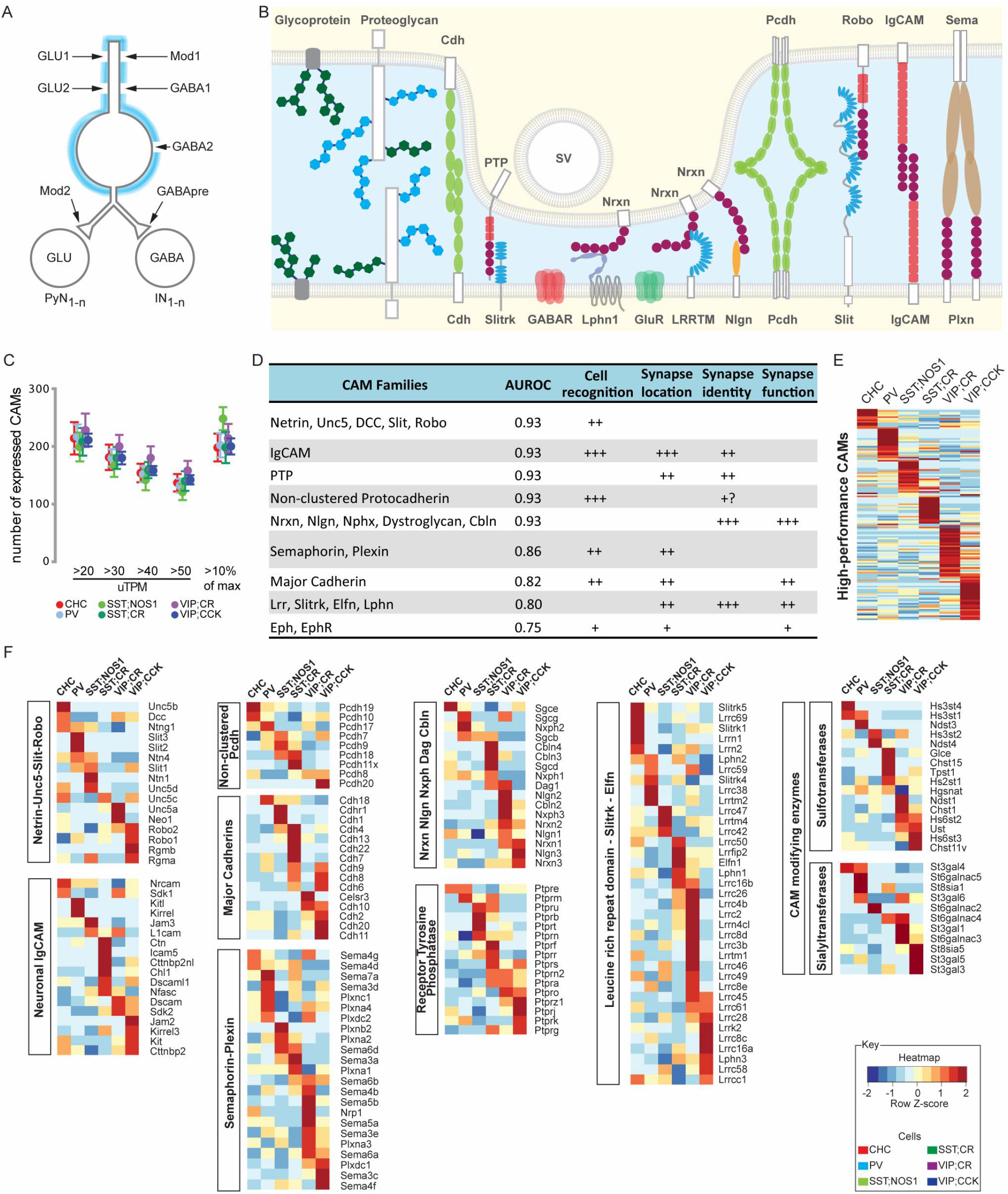
Differential expression of cell adhesion molecules and carbohydrate modifying enzymes among GTPs. **(A)** Schematic representation of the specificity of synaptic connectivity. A single GABAergic neuron receives multiple sources of excitatory (GLU), inhibitory (GABA) and modulatory (Mod) inputs at different subcellular locations and innervates large sets of pyramidal neurons (PyN) and interneurons (IN). Blue shading denotes extracellular matrix. **(B)** Multiple families of cell adhesion molecules, synaptic adhesion molecules, and glycoproteins provide specific extracellular coating, cell surface and synaptic labels. **(C)** Approximately 200 different cell adhesion molecule genes are expressed in each GTPs estimated by a sliding absolute expression value thresholds or by using 10% of maximum expression value as a dynamic cutoff. **(D)** Major categories of ligand-receptor cell adhesion systems and their demonstrated roles in synaptic connectivity. All these adhesion systems are highly discriminative of GTPs as indicated by their high AUROC scores. Number of “+” is an assessment of the degree of involvement in the listed function. **(E)** Heatmap showing 136 cell-adhesion molecules that are differentially expressed across the six GTPs. **(F)** Eight different cell adhesion systems and two families of carbohydrate modifying enzymes (shown in individual heatmaps) are each differentially expressed among GTPs. Abbreviations: SV, synaptic vesicle; Cdh, Cadherin; PTP, protein tyrosine phosphatase; Nrxn, Neurexin; Pcdh, Protocadherin; Robo, Roundabout; IgCAM, Cell adhesion molecules of the immunoglobulin superfamily; Sema, Semaphorin; Slitrk, SLIT And NTRK Like Family Member; GABAR, GABA receptor; Lphn1, Laterophilin1; LRRTM, Leucine Rich Repeat Transmembrane proteins; Nlgn, Neuroligin; Slit, Slit Guidance Ligand; Plxn, Plexin.

For example, the Netrins mediate attractive interaction through DCC receptors and repulsive interaction through UNC5 family members, and the Slits regulate axon branching and mediate repulsive actions through the ROBO receptors (Kolodkin and Tessier-Lavigne, 2011; Wang et al., 1999). We found that different UNC5 members are expressed in ChC, SST/nNOS, SST/CR, VIP/CR cells. Double fluorescence mRNA in situ confirmed that UNC5b is highly specific to ChCs (Figure S2B). Furthermore, Unc5a, 5c, 5d and their ligand netrin1 are differentially expressed among GTPs. These receptor-ligand pairs might mediate cell-cell recognition (e.g. attraction or repulsion) (Figure S3A). On the other hand, Slit2 and 3 are highly enriched in PV cells and might contribute to the exuberant axon terminal branching that characterizes their “basket-like” morphology. In addition, ~20 immunoglobulin cell adhesion molecule (IgCAMs) are differentially expressed (Figure 3F; Supplemental Text 1a) and may contribute to the cellular, subcellular and synaptic specificity among GTPs. Among them, CHL1 is particularly enriched in SST/CR population which includes dendrite-targeting Martinotti cells, consistent with its role in regulating subcellular synapse specificity (Ango et al., 2008).

Among synaptic adhesion molecules, the neurexin (NRXs) and neuroligin (NLGs) are key pre- and post-synaptic organizers and regulate synaptic assembly and transmission properties through interaction with their associated proteins (Sudhof, 2008). Protein tyrosine phosphatases (PTPs) represent another crucial set of presynaptic organizers (Takahashi and Craig, 2013). On the postsynaptic side, a large family of leucine rich repeat proteins (LRRs), including LRR transmembrane proteins (LRRTMs) and Slitrks, interact with presynaptic RPTPs and NRXs to regulate synapse diversity, specificity and plasticity (de Wit and Ghosh, 2014). We found that each of these synaptic adhesion families is differentially expressed among GTPs (Figure 3F). Notably, each GTP enriches for a different set of 6-12 LRR proteins (Figure 3F). Among these, Elfn1 is prominently enriched in SST/CR cells (Figure 3F). Elfn1 is also enriched hippocampal O-LM interneurons - a homologue of cortical Martinotti cells contained within the SST/CR population, and contributes to the synaptic facilitation of glutamatergic transmission onto O-LM cell (Sylwestrak and Ghosh, 2012), a property also shared by Martinotti cells (Silberberg and Markram, 2007). Cell specific expression of LRRs might contribute to post- and trans-synaptic specializations that customize the property of synapse types defined by pre- and post-synaptic neuron identities.

We further discovered prominent differential expression in two families of carbohydrate modifying enzymes that may increase the molecular diversity of glycosylated CAMs and proteoglycans on cell membrane and in extracellular matrix (Figure 3F; Figure S3B; see Supplemental Text 1b).

Together, our results suggest that transcription profiles of GABAergic neurons encode molecular mechanisms to diversify and specify not only their cell membrane but also extracellular milieu. Each GTP might produce a characteristic cell coat through distinct carbohydrate modification patterns to diversify proteoglycans that facilitates or prevents cell interaction at a distance. Further, nearly all families of adhesion molecules that regulate circuit development maintain expression in mature neurons, and almost every family shows substantial differential expression among GTPs. These adhesion families likely constitute a comprehensive mosaic of multi-faceted cell surface code throughout the neuronal membrane. Cell specific alternative mRNA splicing and post-translational glycosylation through carbohydrates and sulfation patterns will further increase the diversity, specificity and flexibility of this cell surface code.

### Differential expression of transmitter and modulator receptors shapes input properties of GTPs

Cortical GABAergic neurons received a large variety of extracellular inputs mediated by neurotransmitters, modulators, hormones and cell contacts that exert their actions through three broad classes of surface receptors: ligand-gated ion channels, G-protein coupled receptors, and enzyme-coupled receptors (e.g. receptor tyrosine kinases and phosphatases). Each class contains dozens to hundreds of receptors encoded by multiple gene families (Luo, 2016). Each receptor is characterized by unique ligand binding specificity, biophysical and biochemical properties, signaling properties, and subcellular localization. This broad receptor repertoire endows neurons with the large capacity to detect and transduce multiple extracellular signals with appropriate specificity and flexibility. We found that nearly every receptor family in each broad class is differentially expressed among GTPs (Figure 4).

**Figure 4:**
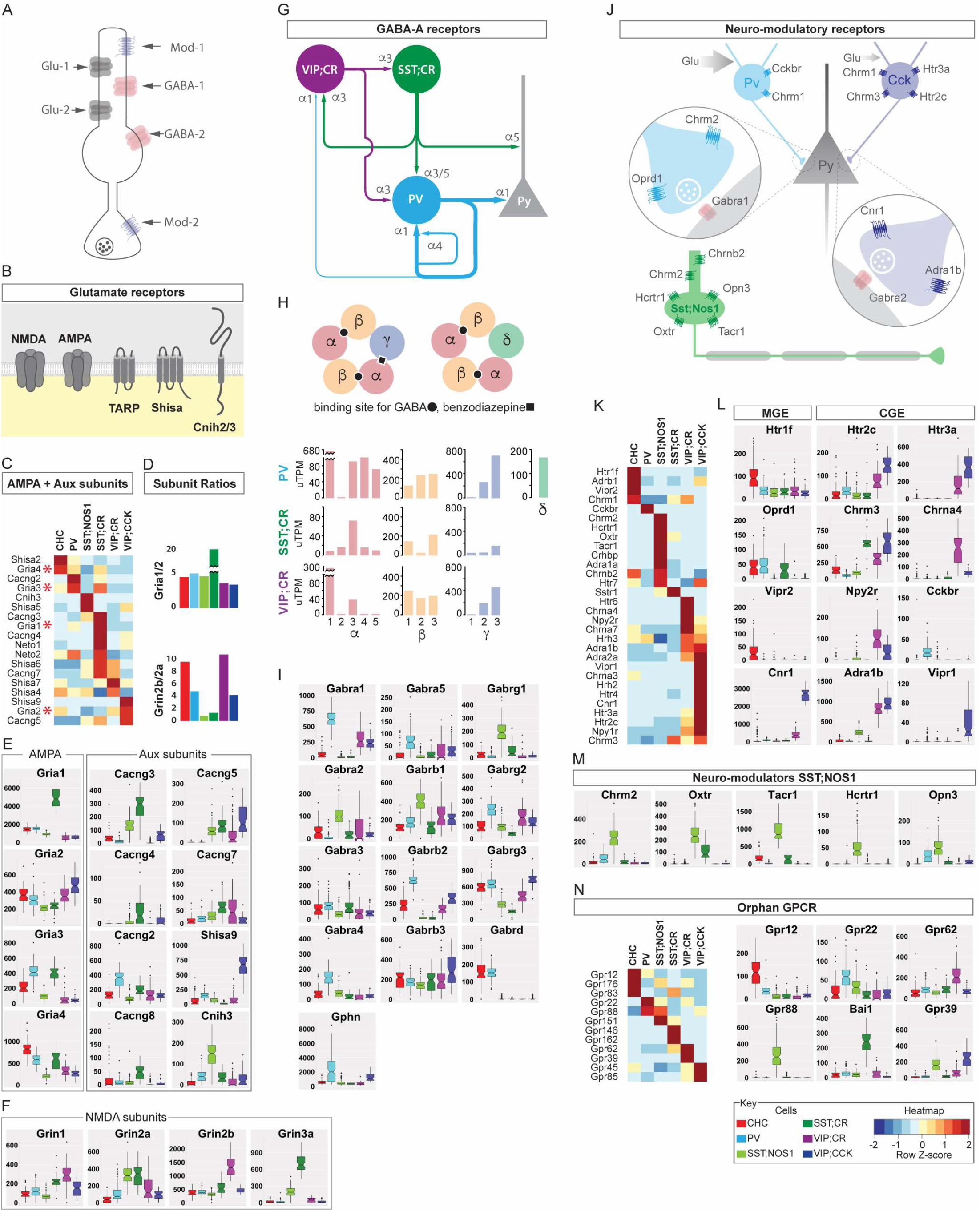
Differential expression of transmitters and modulatory receptors among GTPs. **(A)** Schematic of various excitatory, inhibitory and modulatory receptors expressed on a generic GABAergic neuron. **(B)** Schematic of glutamate receptor core subunits and auxiliary proteins that together form native receptors with specific localization, trafficking and biophysical properties. **(C)** Heatmap showing differential expression of AMPAR core subunits and auxiliary proteins across GTPs; SST;CR cells express the greatest diversity of AMPARs. **(D)** Top: SST;CR cells show highest Gria1 (GluA1)/Gria2(GluA2) ratio among GTPs. Bottom: Most GABAergic neurons have more Grin2b (GluN2B) than Grin2a (GluN2A) receptors but the reverse is true in SST neurons. **(E)** Select boxplots of AMPAR core and auxiliary subunits shows striking level differences among GTPs; y-axis in uTPM counts. **(F)** Select boxplots of NMDA subunits; glycine-activated Grin3a (GluN3A) is highly expressed in SST;CR cells. **(G)** Schematic summary of GABA_A_R subunit expression among GTPs deduced from combining transcriptome analysis and literature information. Note that particular GABA_A_R subtypes may match specific types of presynaptic GTP terminals. PV and SST/CR cells, respectively, have the most and least diverse GABA_A_Rs and inhibitory inputs. **(H)** Top: Schematic of the pentameric subunit composition of GABA_A_R and ligand binding sites. Bottom: Differential expression of α, β and γ subunits and the stoichiometric constraints of their pentameric combinations determine the possible diversity of GABA_A_R subtypes within a GTP; note that PV and SST/CR cells have the most and least diverse diversity, respectively. **(I)** Select boxplots showing subunit level differences among GTPs. PV cells have the highest levels of α1, α4, α5 and also the inhibitory postsynaptic scaffolding protein Gphn (Gephrin). **(J)** Schematic comparison of the neuromodulatory receptors among PV and CCK basket cells and SST;NOS1 long projecting neurons. **(K)** Heatmap of neuromodulatory receptors showing differential expression among GTPs; SST;NOS1 and VIP;CCK cells shows the highest diversity. **(L)** Select boxplot showing that CGE-derived interneurons tend to express more neuromodulatory receptors types compared to MGE-derived interneurons. **(M)** Select boxplots of neuromodulatory receptors specific to or enriched in SST;NOS1 cells. **(N)** Differential expression of orphan GPCRs among GTPs shown as heatmap (left) and boxplots (right).

### Ionotropic glutamate receptors (iGluRs)

Ionotropic glutamate receptors (iGluRs) play key roles in excitatory synaptic signaling and plasticity and include: AMPA (GluA1–4), NMDA (GluN1, GluN2A–D, GluN3A–B), and kainite (GluK1–5) receptors (Traynelis et al., 2010). The basic biophysical properties of iGluRs are determined by their tetrameric pore-forming subunits, shaped by subunit composition, alternative splicing and RNA editing. Despite progress in understanding the role of iGluR in well-characterized principle neurons (e.g. CA1 pyramidal neurons) (Huganir and Nicoll, 2013), the picture in GABAergic neurons is far less clear, largely due to the diversity of cell types with distinct properties of glutamate transmission and heterogeneous patterns of iGluR expression (Akgul and McBain, 2016; Moreau and Kullmann, 2013). Here we provide quantitative mRNA profiles of iGluRs and auxiliary subunits in GTPs (Figure 4B-E), which suggests the potential for cell type specific assembly of a large variety of native AMPARs with customized distribution patterns and functional properties.

Glutamatergic synapses in GABAergic interneurons often contain higher proportions of CP-AMPARs (Jonas et al., 1994; McBain and Dingledine, 1993) and GluN2B-NMDARs (Lei and McBain, 2002), although the ratio between the two types of AMPARs and NMDARs vary significantly among different cell populations(Akgul and McBain, 2016). Consistent with and substantiating previous physiological results largely from hippocampal interneurons (Akgul and McBain, 2016), we found that the mRNA levels and relative ratio of CP-vs CI-AMPAR subunits in PCPs vary in a highly cell type-dependent pattern (Figure 4B-D). CGE-derived VIP cells have overall relatively low AMPARs and roughly similar GluA1 and GluA2 levels (GluA1:GluA2 = 1.4), and VIP/CR cells have relatively more NMDARs especially those containing GluN2B (GluN2B:GluN2A = 11.0). On the other hand, MGE-derived cells have much higher levels of GluA1 (average GluA1:GluA2 = 8.4), with striking cell type differences: GluA1:GluA2 ranges from 4.1 in SST/NOS1 cells to 20.4 in SST/CR cells (Figure 4D). While PV cells have highest GluA3 levels and CHCs have highest GluA4 levels, SST/CR cells show highest levels of GluA1 and highest non-GluA2/GluA2 ratio (24.8). Interestingly, SST/CR cells also have relatively high GluN2A:GluN2B ratio for NMDARs among the PCPs (Figure 4D). These results suggest cell type-dependent composition and correlation of AMPA and NMDA receptor pore-subunits, especially with regard to the relative abundance and ratio of CP-vs CI-AMPARs and 2B-vs 2A-NMDARs.

In addition to the pore-forming subunits, native AMPARs incorporate multiple auxiliary subunits that regulate AMPAR membrane trafficking, synaptic targeting, gating and signaling (Haering et al., 2014; Jackson and Nicoll, 2011; Straub and Tomita, 2012). The large number and multiple families of AMPAR auxiliary proteins and their regional and cell type specific expression suggest that differential combinations of pore-forming and auxiliary subunits may assemble a large variety of native AMPARs with distinct synaptic distribution patterns and biophysical properties (Dawe et al., 2016; Khodosevich et al., 2014; Tao et al., 2013), but the expression patterns of these auxiliary subunits in GABAergic neurons are largely unknown. Our transcriptome analysis revealed that TARP, SHISA and CNIH family auxiliary subunits show striking cell specific expression patterns (Figure 4C-E). TARPγ2 is enriched in PV cells, TARPγ3, γ8 and SHISA6 are enriched in SST/CR cells, TARPγ3 and SHISA9 are enriched in VIP/CCK cells. While PV cells predominantly express one auxiliary subunit (TARPγ2), SST/CR cells express at least 6 types (TARPγ2, γ3, γ8, γ5, γ7, SHISA6). Whereas pore-subunits differ in expression levels, auxiliary subunits often show ON/OFF expression among GTPs (Figure 4E). These results suggest that different GABAergic neurons may assemble a specific set of native AMPARs with distinct pore and auxiliary subunit compositions, postsynaptic distribution patterns and biophysical properties. This large repertoire of native AMPARs may achieve cell type- and synapse-specific transmission and plasticity of glutamatergic inputs according to different presynaptic sources.

Taken together, these results suggest that, instead of receiving a more or less generic set of glutamatergic inputs, different GABAergic neurons likely deploy a distinct set of native AMPARs to customize the amplitude, duration, dynamics, and thus the shape of glutamate synaptic currents in a cell and synapse specific manner. (See summary in Table 1 for the striking case of SST/CR cells).

**Table 1.**
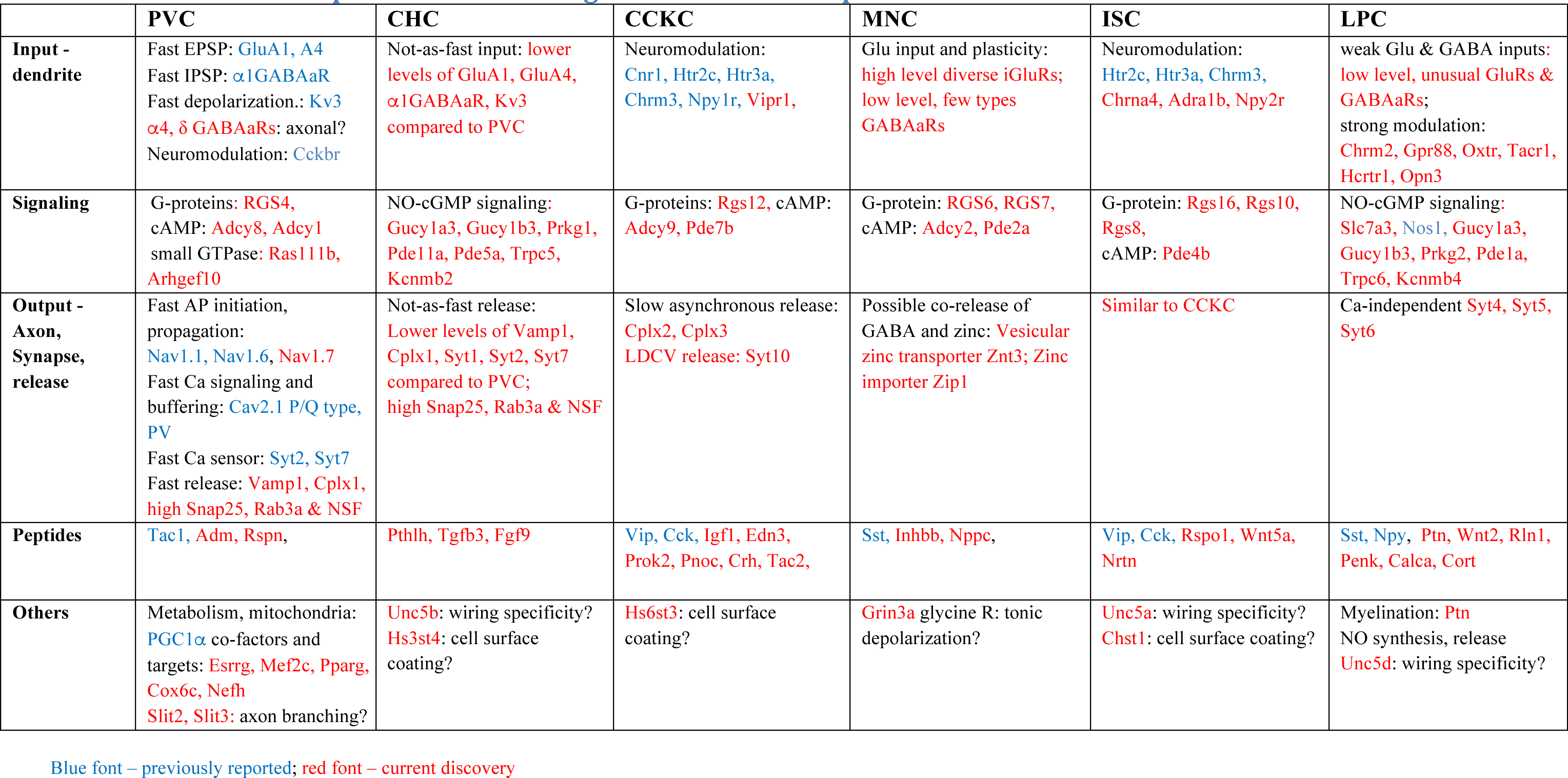
Molecular portraits of GABAergic Ground Truth Populations.

### Ionotropic GABA receptors (GABA_A_Rs)

GABA_A_ receptors mediate fast inhibitory neurotransmission and are assembled as heteropentameric chloride channels, typically consisting of 2α, 2β, and 1γ subunits (Olsen and Sieghart, 2008). The subunit composition critically determines their kinetics, pharmacology, and subcellular distribution. Over a dozen of the total 19 subunits are expressed in the brain (e.g. α1-6, β1-3, γ1-3, and δ), suggesting the combinatorial potential for a very large number of GABA_A_R subtypes, but subunit partnership is thought to be governed by preferential assembly to form a more limited number of subtypes. Although multiple GABA_A_R types have been demonstrated in *in vitro* expression systems, to date only a dozen native GABA_A_Rs with known subunit composition have been identified based on co-expression, electrophysiological and pharmacological evidence (Olsen and Sieghart, 2009). The vast majority of possible subunit combinations remain tentative, in part because most studies only achieved brain regional but not cellular resolution of subunit expression and co-expression. Here we provide comprehensive and quantitative mRNA profiles of all GABA_A_R subunits in GTPs, which reveal highly cell type specific repertoire of GABA_A_R subtypes.

Whereas γ2 is ubiquitously present in all neurons and regarded as the obligatory subunit of most if not all synaptic GABAaRs that mediate phasic inhibition, γ3 is sparsely expressed in cortical neurons of unknown identity (Olsen and Sieghart, 2008). Although γ3 can assemble with α and β to form synaptic receptors with slowly decaying IPSCs (Kerti-Szigeti et al., 2014), its cellular expression and physiological significance is unclear. We found that, surprisingly, γ3 is not only prevalent but also transcribed at much higher levels than γ2 subunits in all 6 GTPs (Figure 4H). This suggests that γ3 may uniquely contribute to the assembly of a class of slower decaying, longer duration synaptic GABA_A_Rs in GABAergic neurons.

Furthermore, different GTPs show highly specific subunit profiles and levels (Figure 4H, I). PV cells express the largest variety (all except α2, α6, γ1) and overall highest levels of subunits, and uniquely high level of the GABA_A_R clustering/scaffolding protein gephyrin. In contrast, SST/CR cells express the least variety (mainly α3, β1/3, γ2/3) and lowest overall levels. Interestingly, SST/nNOS cells are distinguished by predominant expression of slow kinetics α2-containing GABA_A_Rs and, surprisingly, the exceedingly rare γ1 subunit which is thought to assemble extra- or non-synaptic GABAaRs (Dixon et al., 2014). On the other hand, PV and ChCs express the δ subunit, known to assemble extra-synaptic GABAaRs (especially with in combination with α4 – highly enriched in PV cells) that localize to presynaptic terminals (Belelli et al., 2009; Herd et al., 2013).

These comprehensive and quantitative cell resolution profiles, when considered together with the well-characterized connectivity patterns among GTPs, suggest that distinct GABA_A_R subtypes with specific subunit combinations are likely targeted to specific connections that match the presynaptic terminals to optimize inhibitory transmission properties (Figure 4G). For example, PV cells predominantly mediate self-inhibition (i.e. other PV cells) in addition to perisomatic inhibition of pyramidal neurons (Jiang et al., 2015; Pfeffer et al., 2013). They further receive inhibitory inputs from SST cells (mainly SST/CR positive Martinotti cells) and interneuron-selective VIP cells (mostly VIP/CR positive). Importantly, PV-PV transmission is one of the fastest in the brain, mediated by the α1β2γ2 subtype (Hu et al., 2014; Klausberger et al., 2002). Based on these considerations, it can be inferred that γ3-containing slow kinetics receptors, likely abundant in PV cells, are excluded from PV-PV synapses. Further, the unique co-expression α4 and δ, a well-established combination for extrasynaptic and axonal GABA_A_R, suggest the presence of this subtype in PV cell terminals, likely activated by GABA spill-over (Herd et al., 2014) during concerted GABA release from dense perisomatic synapses characteristic to PV axon terminals. These considerations further raise the possibility that other subunit combinations, such as those containing α3, α5, and γ3, might support inputs from SST/CR and VIP/CR cells, especially in the dendritic compartment of PV cells (Ali and Thomson, 2008). Following similar logic, we infer that SST/CR cells receive VIP cell inputs (Jiang et al., 2015; Pfeffer et al., 2013) likely through α3β1/3γ3 type GABA_A_Rs, and VIP cells likely receive PV cell input (Jiang et al., 2015; Staiger et al., 1997) through α1-containing GABA_A_Rs and Martinotti cell input (Jiang et al., 2015; Staiger et al., 1997) through α3-containing GABA_A_Rs (See Supplementary text 2).

Together, our results suggest that cell type specific subunit expression may allow assembly of specific repertoires of GABA_A_R subtypes, which are endowed with distinct biophysical and pharmacological properties, subcellular localization, and are targeted to specific postsynaptic sites that match presynaptic properties. This exquisite synapse specificity of receptor subtypes might customize inhibitory transmission properties between specific cell types (Figure 4G).

### Neuromodulatory and G-protein coupled receptors

Cortical GABAergic neurons received a wide range of subcortical modulatory inputs that convey diverse signals of brain and behavioral states. These modulators, peptides and hormones act through a large family of G-protein coupled receptors (GPCRs) (Davenport et al., 2013), which trigger multiple signaling pathways that modulate ion channel properties and regulate electrical signaling and transmitter release (Luo, 2016). Decades of studies have revealed cell specific expression and function of neuromodulatory receptors in hippocampal interneurons (Armstrong and Soltesz, 2012), but a comprehensive picture of modulatory receptors across cell types have not been achieved. Here we present comprehensive and quantitative transcription profiles of neuromodulatory receptors in GTPs.

Whereas MGE-derived interneurons (ChC, PV, SST/CR cells) are characterized by higher levels and larger variety of iGluRs and GABA_A_Rs, CGE-derived interneurons express much larger variety of neuromodulatory receptors (Figure 4K-L). This broad distinction is best illustrated by a comparison of PV and VIP/CCK cells (Figure 4J-L), both innervate the perisomaitc regions of pyramidal neurons and are extensively studied in hippocampal CA1 (Armstrong and Soltesz, 2012; Freund and Katona, 2007). We confirmed most findings derived from the hippocampus: whereas PV cells show enrichment of only a few modulatory receptors (e.g. CCK2R, Oprd1), VIP/CCK cells express multiple GPCRs for serotonin, acetylcholine, norepinephrin, endocannabinoid. We further discovered that VIP/CCK cells also express Adra1b, NPY and VIP receptors. Considered together with their iGluR and GABA_A_R profiles, these results suggest that, similar to their homologs in the hippocampus, cortical PV and CCK basket cells represent two highly distinct cell types that likely provide different “flavors” of perisomatic inhibition (Freund and Katona, 2007): while the former is recruited by fast and precise excitatory and inhibitory inputs from local and cortical sources, the latter is profoundly modulated by subcortical inputs that represent mood, internal drive, and behavioral state.

As a clear exception among MGE-derived GABA neurons, the unique long axon projection of SST/nNOS cells is associated with multiple unusual features, including neuromodulatory inputs. Contrasting other MGE cells, SST/nNOS cells express lower levels of iGluRs and extra- or non-synaptic γ1-containing GABA_A_Rs. On the other hand, they express a large and unusual set of modulatory receptors including hypocretin, oxytocin, neurokinin, Tacr1 (Figure 4M), which are released from hypothalamic centers that regulate global brain states (Kilduff et al., 2011; Schwartz et al., 2016). Together, these results depict a cell type with weak phasic excitatory and inhibitory inputs but a wide range of tonic subcortical modulatory inputs, consistent with its activation by homeostatic sleep drive, and speculated role in regulating global cortical networks (Kilduff et al., 2011; Schwartz et al., 2016.

GTPs are further characterized by their expression of orphan GPCRs for unknown or unproven ligands. Each GTP can be distinguished from all other by unique or highly enriched expression of at least 2 orphan GPCRs (Figure 4N). Although the function of most these GPCRs are unknown, the metabotropic Zn^2+^ sensor GPR39/mZnR (Perez-Rosello et al., 2013) is specifically expressed in VIP/CCK and to a less extent SST/nNOS cells. Recent studies suggest that, upon Zn^2+^ binding, which is co-released with glutamate and possibly other transmitters, GPR39 promotes KCC2 membrane trafficking, thereby enhancing GABA_A_R mediated hyperpolarization (Chorin et al., 2011). Thus GPR39 in VIP/CCR cell might mediate activity-dependent modulation of their excitability.

In summary, cell type profiles of modulatory receptors are highly congruent with their ionotropic receptor profiles and together may support the distinct recruitment and modulatory properties of each cell type. Cell specific repertoire of nearly all major families of ligand-gated receptors may endow GABAergic neuron the capacity to detect and transduce specific combinations of transmitters and modulators in a characteristic manner to elicit appropriate responses.

### Differential expression of voltage-gated ion channels and electrophysiological properties of GTPs

GABAergic neurons maintain their characteristic ionic balance to shape intrinsic membrane potential and firing properties. They respond to synaptic and modulatory inputs with changes in local membrane potentials that integrate and initiate action potentials, which propagate to axon terminals and trigger transmitter release and other physiological responses. These highly sophisticated electrophysiology properties and ion homeostasis are shaped by several families of voltage-gated ion channels (VGICs), each contains diverse family members with characteristic biophysical properties (Yu and Catterall, 2004). Among these, sodium channels (Nav) drive the initiation and propagation of membrane depolarization and action potential (Kruger and Isom, 2016), potassium channels (Kv) regulate membrane re-polarization (Trimmer, 2015), and Calcium channels (Cav) transduce membrane potential changes into intracellular Ca2+ transients that initiate many physiological events (Zamponi et al., 2015). In addition to voltage dependence, intracellular signals (e.g. Ca^2+^, H^+^, ATP, cyclic nucleotides) regulate calcium-activated (Kca), inward rectifier (Kir), and 2-pore (K2p) potassium channels (Trimmer, 2015). The ion selectivity, gating and regulation of these channels are tailored to shape specific aspects of electrical signaling. These channels are further targeted to subcellular compartments, often regulated by auxiliary subunits and linked to customized signaling complexes, to optimize electrical signaling at designated microdomains (Dolphin, 2016; Kruger and Isom, 2016; Vacher et al., 2008; Vacher and Trimmer, 2011). Except in rare cases (Hu et al., 2014), comprehensive profiles of VGICs in specific neuronal subpopulations have not been described. Our transcriptome analyses demonstrate extensive differential transcription profiles within and across multiple VGICs families among GTPs (Figure S4).

Within the Nav and Cav family, major pore-forming subunits are broadly expressed among GTPs, often with different expression levels (Figure S4C-E). Interestingly, Cav auxiliary subunits (β1-2, α2, δ1-4) show more distinct, often binary (ON/OFF) pattern (Figure S4D), suggesting cell specific regulation of the trafficking, gating, and kinetics of pore forming subunits. Within the Kv family, different gene subsets are prominently enriched in each of the GTPs (Figure S4C). Importantly, there is a tight correlation between the expression of Kv principle subunits (e.g. Kcna1/Kv1.1 and Kcna2/Kv1.2) and their matching auxiliary subunits (Kvβ1-3, Kcnab1, b2 & b3) in specific GTPs (e.g. PV cells), suggesting cell specific assembly of functional channel complex.

Although mRNA levels do not linearly translate into protein levels and their subcellular distribution patterns, the relevance of ion channel transcription profiles to physiological properties is highlighted by the striking case of PV cells. Fast-spiking PV cells convert an excitatory input signal to an inhibitory output signal within a millisecond, a stunning cell biology feat that appears to involve optimizing multiple aspects of electrical signaling across subcellular compartments, in part through specific expression and localization of a unique assortment of VGICs with highly tailored biophysical properties (Hu et al., 2014). For example, Kv1 and Kv3 promote short action potential duration and sublinear summation, Nav1.1 (Scn1a) and Nav1.6 (Scn8a) facilitate fast AP initiation and propagation, and Cav2.1 (P/Q) promotes fast GABA release (Hu et al., 2014). Our PV cell transcription profile confirms, and thus is validated by, these published molecular and electrophysiological studies. Our results further allow quantitative comparison of each channel gene expression across GTPs. For example, the prominent enrichment of multiple Navs (Scn9a, Scn8a, Scn1a, Scn3a, Scn1b, Scn4b, Scn2b) likely underlie the “supercritical density” of Nav for ensuring fast signaling in PV cell axons (Hu and Jonas, 2014), and the striking elevation of a large set of fast-kinetics Kv1-4 members may implement rapid repolarization and narrow AP duration at each subcellular domain. In addition, our results reveal novel expression in auxiliary subunits Kca, Kir and K2p, which hint other uncharacterized physiological properties. These findings suggest that ion channel transcription profiles in other less characterized GTPs may similarly predict physiological features that can be validated by experimental studies. Together, our results suggest that differential and correlated expression across multiple families of VGIC subunits may customize the electrical signaling among GTPs.

### Differential expression of signaling proteins in calcium, cyclic nucleotide and small GTPase 2^nd^ messenger pathways customizes intracellular signaling in GTPs

In addition to fast electrophysiological responses, extracellular signals trigger a variety of metabolic, morphologic, transcriptional and neurosecretory responses. The conversion of specific combination of inputs to a concerted set of short-term physiological and long-term adaptive responses is mediated by myriad intracellular signaling pathways. As a universally conserved cell signaling scheme (Alberts, 2014), a large repertoire of surface receptors transduce diverse extracellular signals into a small set of intracellular 2^nd^ messengers such as Ca^2+^, cyclic nucleotides (e.g. cAMP, cGMP), lipid metabolites (e.g. diacylglycerol) and small GTPases (e.g. Ras, Rho); these 2^nd^ messengers typically trigger enzyme cascades that engage different sets of effector proteins to execute cell responses in excitability, transmitter release, metabolic rate, neurite motility and gene expression (Figure 5A) (Luo, 2016). Studies from mostly non-neuronal systems have demonstrated that, superimposed upon several highly conserved schemes of 2^nd^ messenger cascades, different cell types deploy a large set of regulatory signaling proteins to control the spatiotemporal dynamics of each 2^nd^ messenger and signal transduction to specific effector systems to achieve appropriate cell responses (Brini et al., 2014; Halls and Cooper, 2011; McCormick and Baillie, 2014). The mammalian genome contains dozens of gene families that encode hundreds of signaling proteins associated with just a handful of major 2^nd^ messenger systems (Alberts, 2014). Whether and how different neuron types coordinate the expression and action of multiple families of signaling proteins to customize signal transduction that translates specific input to appropriate output is almost entirely unknown. Through our computation screen of gene families, we discovered that, whereas most kinase cascades and signal proteins are broadly expressed, a small set of regulatory protein families in the calcium, cyclic nucleotide and small GTPase pathways are highly differential among GTPs and may tailor specific properties of their signal transduction (Figure 5).

**Figure 5:**
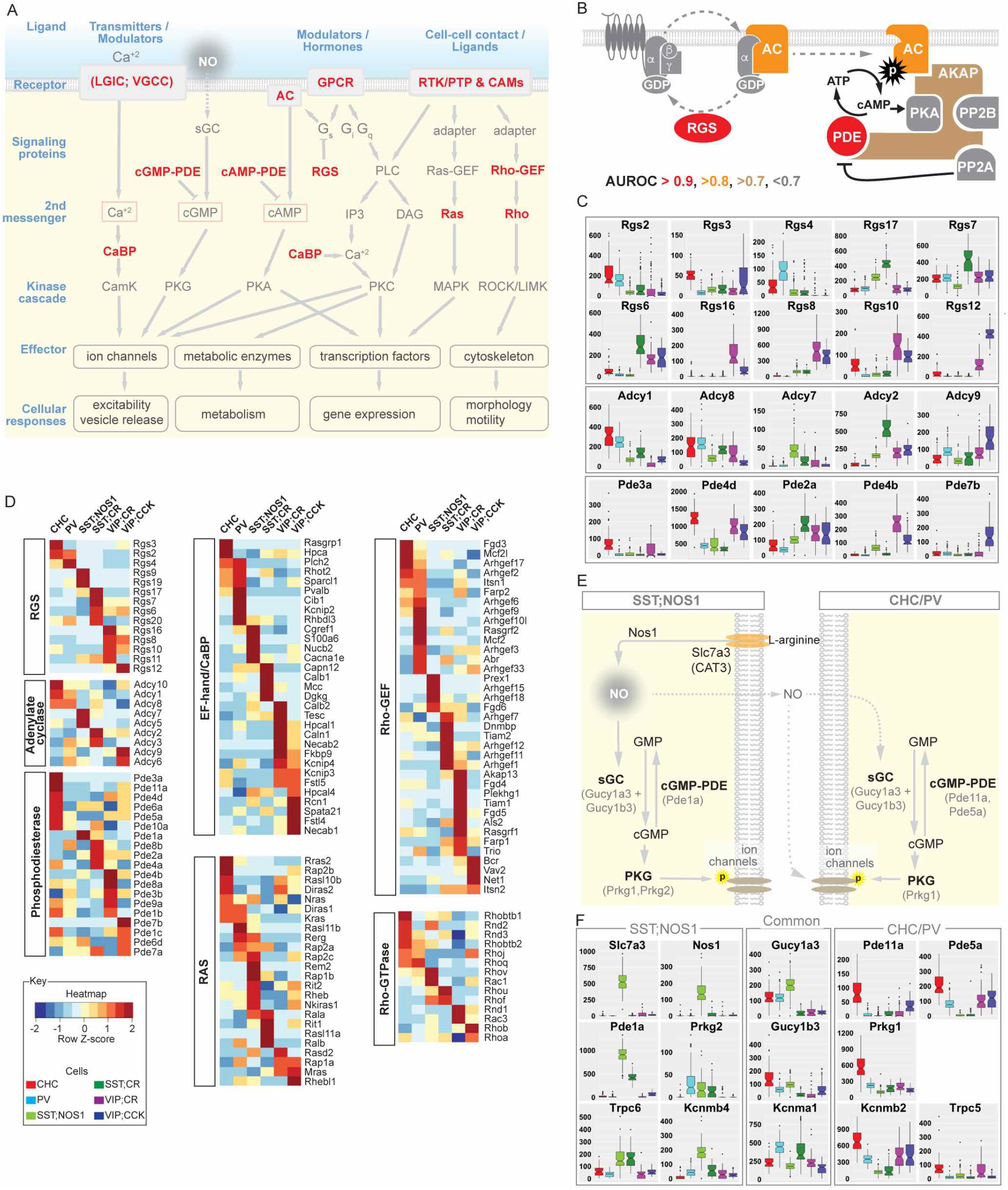
Differential expression of signaling proteins in several 2^nd^ messenger pathways customizes intracellular signaling in GTPs. **(A)** A simplified schematic summary showing that the Ca, cAMP, cGMP, Ras and Rho signaling pathways are differentially configured among GTPs. While the core skeletons of signal transduction machineries, kinase cascades and effectors are common among GTPs (grey, with low AUROC scores), a small set of regulatory signaling proteins (red) are strikingly differentially expressed with high AUROC values. **(B)** Schematic of a GPCR signaling module illustrating that while multiple components (grey) are common among GTPs, different members of key regulatory proteins such as RGS, Adenylate cyclases (AC), Phosphodiesterases (PDE) and A-kinase anchoring proteins (AKAPs) are differentially expressed and likely customize the specificity and spatiotemporal dynamics of cAMP signaling. **(C)** Boxplots showing that different combinations of RGS, AC and PDE members are enriched in individual GTPs. **(D)** Heatmap of several classes of signaling proteins with high AUROC scores shown in 5A. **(E)** Enhanced NO-cGMP signaling in SST;NOS1 and CHC cells. The entire pathway of NO synthesis and cGMP production (guanylyl cyclase), degradation (PDE), kinase signaling (PKG) and putative phosphorylation targets are coherently and specifically expressed or enriched in SST;NOS1 cells as well as CHC cells. **(F)** Boxplots show differential expression in key components of NO-cGMP signaling (depicted in 5E) among GTPs; note ON/OFF patterns or dramatic level differences.

### Ca^2+^ binding proteins likely shapes spatiotemporal dynamics of Ca^2+^ signaling

We found that each GTP expresses a set of ~5-8 different Ca^2+^-binding proteins (CaBPs; Figure 5D). Many of these CaBPs are in fact signaling proteins (e.g. Rasgrp1 in ChCs). These results suggest that differential expression of multiple Ca^2+^ binding and signaling proteins might shape distinct spatiotemporal dynamics and the specificity of Ca^2+^signaling among GTPs (see Supplemental Text 3a).

### Adenylyl cyclase and phosphodiesterase members may shape distinct cAMP signaling properties

GPCRs signal through G proteins, many of which engage cAMP - the archetypical 2^nd^ messenger pathway. cAMP activates protein kinase A (PKA) which regulates effector proteins through phosphorylation (Supplemental Text 3b). The synthesis, degradation and spatiotemporal dynamics of cAMP are stringently regulated at each step (Halls and Cooper, 2011). We found that, while the G protein subunits themselves are broadly expressed, regulators of G protein signaling (RGS; (Gerber et al., 2016) family members manifest highly differential expression, often with binary-ON/OFF patterns among GTPs (AUR0C=0.93; Figure 5A-C); this suggests that the turning-off of G_α_ subunit, a crucial step of G protein regulation, is implemented in a cell specific manner. Downstream to G proteins,7 of the 9 adenylyl cyclases (ACs) members with different catalytic and regulatory properties are differentially expressed (AUROC=0.85): whereas the Ca^2+^/Calmodulin-activated AC1 and AC8 (Halls and Cooper, 2011) are enriched in PV and ChC cells, the PKC-activated AC7 and AC2 (Halls and Cooper, 2011) are enriched in SST/NOS1 and SST/CR cells, respectively (Figure 5A-C). More strikingly, phosphodiesterases (PDEs), which mediate rapid cAMP degradation (Maurice et al., 2014), is among the top differentially expressed gene families (AUROC=0.94): 15 of the 22 members are differentially expressed, often with ON/OFF patterns. For example, Pde11a, 1a, 4b, 7b are each highly enriched in ChC, SST, VIP/CR, and VIP/CCK populations (Figure 5A-C). Substantial evidence in non-neuronal cells have demonstrated that different PDE members are targeted to highly confined subcellular compartments, in part through recruitment by specific A kinase adaptor proteins (AKAPs) into signaling complexes (Edwards et al., 2012). It has been hypothesized that the assembly of these subcellular targeted “signalosomes” containing particular members of synthetic and degradation enzymes with distinct catalytic and regulatory properties, contributes to both the fine-tuning and specificity of compartmentalized cAMP signaling (Maurice et al., 2014). Although the specific combinations of AC, PDE and AKAP members and their functional effectors in GTPs remain to be elucidated, their specific and correlated transcription patterns suggest possible mechanisms whereby the spatiotemporal patterns of a single ubiquitous 2^nd^ messenger can be crafted to direct receptor-(i.e. input) and cell-specific signal transduction in different GTPs.

### cGMP signaling modules in SST/nNOS and ChC

In contrast to cAMP, which serves as a ubiquitous 2^nd^ messenger for vast number of extracellular ligands through hundreds of GPCRs, cGMP signaling in the brain is predominantly triggered by nitric oxide (NO) (Lucas et al., 2000). In mature cortex, nNOS is expressed in subsets of GABAergic neurons, with high levels in a small set of SST^+^ long projection cells (LPCs, also type I nNOS cells) and much lower levels in several other populations (type II nNOS cells) (Perrenoud et al., 2012; Taniguchi et al., 2011). Although the general scheme of NO signaling is well established in brain tissues (Supplemental Text 3c), whether NO and cGMP signaling is differentially implemented in different neuronal cell types is far from clear. We found that not only the synthetic enzyme nNOS is specific to LPCs, so is the expression of the major neuronal L-arginine transporter Slc7a3 that supplies the substrate for NO synthesis (Figure 5E; (Friebe and Koesling, 2003). This tight co-expression likely contribute to a coordinated mechanism that endows LPCs as the major source of cortical NO and further suggests that type II nNOS neurons not only have low levels of the synthetic enzyme but also low levels of substrate for NO production. As the key link from NO to cGMP production, the soluble guanylyl cyclase (sGC) functions as a strict heterodimer of α and β subunits, and the mouse brain mainly contains Gucy1α2, Gucy1α3, Gucy1β3 (Friebe and Koesling, 2003). We found that while Gucy1α2 is expressed at low levels across GTPs, Gucy1α3 and Gucy1β3 are highly enriched in ChC, PV and LPC cells but are nearly absent in SST/CR and VIP cells (Figure 5E). This result suggests that whereas cGMP signaling is likely prominent in the former three cell types, it is weak in the latter three populations. Consistent with this finding, cGMP-degrading Pde1a, 5a, 11a are also highly enriched in LPCs and ChCs (Figure 5E), which may regulate the spatiotemporal dynamics of cGMP in these cells. Among the two types of cGMP-dependent PKGs, Prkg1 is found in all GTPs but with major enrichment in ChCs (Figure 5E). These results suggest that, unlike cAMP as a truly ubiquitous 2^nd^ messenger, cGMP specializes to mediate NO signaling in specific cell types.

Furthermore, we found at least two members of the transient receptor potential channels (Trpc5, Trpc6; (Takahashi et al., 2008; Yoshida et al., 2006) and BK-type potassium channels (α1 core subunit and β auxiliary subunits of KCNMA1; (Alioua et al., 1998; Kyle et al., 2013; Zhou et al., 2001)) that are differentially enriched in these two cell types (Figure 5F) and have been shown to be NO and PKG targets (See Supplemental Text 3c). Together, these results reveal striking differences in the mode of NO-cGMP signaling across GTPs and identified two distinct signaling modules in LPCs and ChCs. The stunning coordination in the expression of multiple (8-9) genes encoding almost the entire NO-cGMP pathway, from ligand synthesis and 2^nd^ messenger signaling to potential effectors, can hardly be explained without invoking the transcriptional orchestration by an underlying cell type gene regulatory network.

### Differential expression of Ras and Rho small GTPases

In addition to Ca^2+^ and cyclic nucleotides, many cell surface receptors signal through a large set of Ras superfamily small GTPases to activate multiple kinase cascades that engage effectors (Alberts, 2014; Colicelli, 2004). Prominent among these effectors are transcription factors, which regulate gene expression (Ye and Carew, 2010), and cytoskeleton proteins that regulate cell shape, motility, adhesion and intracellular transport (Soderling, 2014). The mammalian genome contains ~30 Ras-GTPases and ~ 20 Rho-GTPases, and each is regulated by several dozens of guanine nucleotide exchange factors (GEFs) and inactivated by GTPase activating proteins (GAPs) (Cherfils and Zeghouf, 2013). Whether Ras and Rho signaling in the brain are tailored to the needs and properties of different neuron types are unknown, in part due to a near absence of knowledge on their cellular expression patterns.

We found that, within the Ras family, 21 of the 32 members showed major enrichment in specific GTPs (AUROC=0.84; also see Supplemental Text 3d). As different Ras family members might be activated by different upstream signals, have different cellular functions, and engage different downstream effectors (Buday and Downward, 2008; Mitin et al., 2005), this result suggests that GTPs might use Ras members to relay distinct external inputs and trigger appropriate transcription programs and other effectors that mediate long term cellular changes. Furthermore, both the Rho-GTPases and Rho-GEFs are differentially expressed. 37 of the 57 Rho-GEFs (AURPC=0.82) and 14 of the 19 Rho-GTPases (AUROC=0.72; also see Supplemental Text 3d) are enriched in specific GTPs (Figure 5D). As different Rho members are often activated by designated GEFs (Cook et al., 2014), our results suggest that differential expression of Rho signaling and regulatory components might provide the mechanism and capacity to maintain the diversity of GABAergic neuron morphology, connectivity, and to support different forms of neurite and synaptic motility and plasticity.

Altogether, our results suggest that, among the vast number of intracellular signaling proteins constituting myriad pathways that transduce major categories of extracellular inputs, a relatively small number encoded by just a few gene families are differentially expressed among GTPs and likely customize signal transduction to the need and properties of cell types. These gene families converge onto a handful of 2^nd^ messenger pathways mediated by calcium, cyclic nucleotides and small GTPases. A major theme is that almost all these gene families act close to the plasma membrane, before the kinase cascades. Superimposed upon the core skeletons of signaling pathways common across GTPs, these key regulatory components likely shape the specificity and spatiotemporal dynamics of broadly defined 2^nd^ messengers that translate specific inputs to appropriate effectors and cellular responses. It is likely that cross talks among these 2^nd^ messenger systems and signaling pathways may further enhance the specificity and flexibility of cell type specific signal transduction.

### Differential expression of neuropeptides and vesicle release machinery shape distinct outputs

The single most important physiological action of a nerve cell is influencing the activity of its target cells through the release of neurochemical substances. Indeed, the connectivity to proper synaptic partners, the reception and integration of diverse inputs, and the elaborate electrical and intracellular signaling all serve the final singular purpose of releasing appropriate neurochemicals in appropriate “styles”. Although the general scheme and principle of neurotransmitter release have been elucidated (Sudhof, 2013), the molecular mechanisms underlying the surprisingly diverse styles of vesicular release, which differentially impact postsynaptic responses and circuit operation (Markram et al., 2015), are not well understood. Through MetaNeighbour screen, we discovered a surprising diversity of neurochemical contents among GTPs and correlated differential expression of components of vesicular release machinery that may contribute to different release styles (Figure 6).

**Figures 6:**
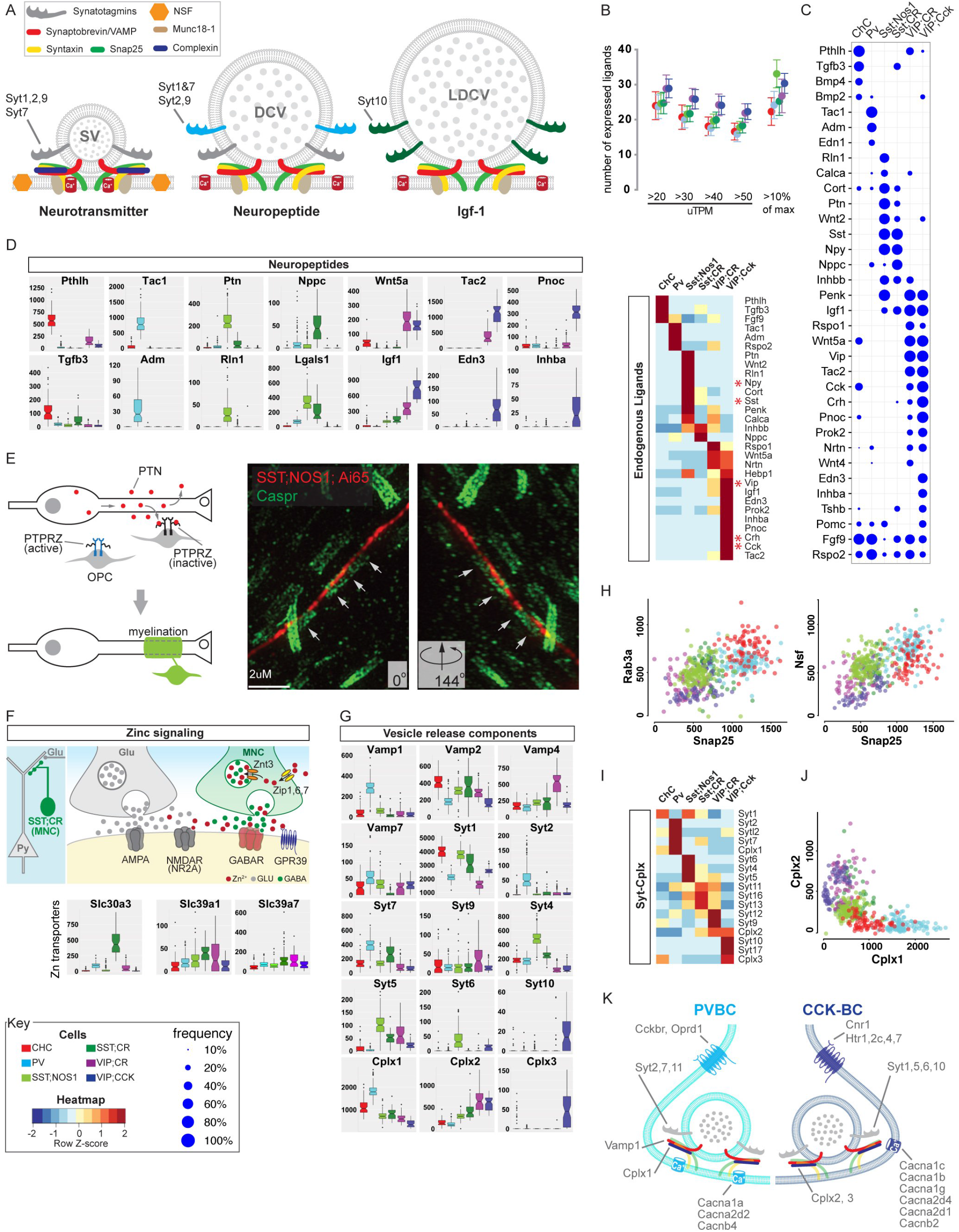
Differential expression of neuropeptides and vesicle release machinery shape distinct outputs and release styles in GTPs. **(A)** Schematic of vesicular release machinery for synaptic vesicle (SV for neurotransmitters), dense core vesicle (DCV for neuropeptides) and large dense core vesicle (LDCV for protein hormones). Putative Syt members implicated in these release machines are listed. **(B)** Top: Each GTP is estimated to express between 20-30 peptides (both unique and common) based on either a sliding threshold of transcript counts (20-50 uTPMs) or dynamic threshold (10% of maximum expression value). Bottom: heatmap shows highly differential expression of endogenous ligands that constitute a neuropeptide code for GTPs. **(C)** Bubble-plot of fraction of individual cells expressing the most commonly expressed neuropeptides among GTPs. Size of dots represents fraction (see key at bottom left). **(D)** Select boxplots showing highly distinct ON/OFF expression of specific neuropeptides in each GTP. Endogenous Ligands is the top gene family (AUROC=0.96) that best distinguishes GTPs. **(E)** Left: Schematic of role of PTN in promoting differentiation of oligodendrocyte precursors for axonal myelination. PTN expression in SST;NOS1 long projection cells predict myelination of their axons similar to excitatory projection neurons. Right: Confirmation of myelination of SST;NOS1 cell axon. 3D render of super-resolution images shows a deep layer RFP-expressing SST;NOS1 axon myelinated by CASPR-labeled oligodendrocyte processes. SST;NOS1 axons are immune-labeled with anti-tdTomato antibody (red) in Sst-Flp;Nos1-CreER; Ai65 animals and co-labeled with anti-CASPR (green). Angular rotation shows coaxial apposition of SST;NOS1 axon and CASPR validating the myelination (white arrows). See Fig-S6 for expanded angular views. **(F)** Schematic showing that Zn may be co-released with GABA specifically from SST;CR terminals. While GABA acts on GABA_A_Rs, Zn may act on nearly non-synaptic NMDARs and influence glutamatergic transmission. Boxplots show high level and specific expression of the Zn vesicular transporter Slc30a3 (Znt3) in SST;CR cells, which also contain the Zn uptake importers Slc39a1 and Slc39a7 (Zip1, Zip7) **(G)** Differential expression of vesicle release machinery components suggest different release styles in Ca^2+^ sensitivity and dynamics among GTPs. **(H)** Scatter plots of mRNA levels (uTPMs) of Snap25 vs Rab3a (left) and Snap25 vs NSF (right). **(I)** Heatmap of Synaptotagmin and Complexin gene families shows selective expression in GTPs. **(J)** Scatterplot of Cplx1 Vs Cplx2 levels (uTPMs) show that fast-release synapses of PV and CHC are biased towards Cplx1 whereas slow-release synapses of VIP;CCK cells mainly utilize Cplx2. **(K)** Comparison summary diagram of PV and VIP;CCK basket cells with contrasting GABA release styles provides molecular correlates of fast-synchronous and slow-sustained vesicle release mechanisms.

### A neuropeptide code of GABAergic neurons

The synthesis and release of different transmitters, peptides and hormones represent a fundamental distinction among neuron types as they produce categorically different outputs that activate different receptors and elicit distinct physiological actions in target cells. However, it is unknown how many peptides are expressed by a GABAergic cell or cell type, and whether peptide expression serendipitously coincide with broad populations or tightly correlates with cell types defined by multiple other features. Our transcriptome analysis revealed a neuropeptide code of GABAergic neurons. We found that over 40 neuropeptides, hormones and secreted ligands are expressed in over 50% of single cells of the 6 GTPs, and each GTP expresses ~3-10 different endogenous ligands (Figure 6B). Single cell analyses demonstrate that individual neurons express multiple peptide and protein ligands (Figure 6C). Importantly, differential expression of these ligands is the most discriminating gene family for GTPs (AUROC=0.96). Indeed, multiple GTPs are uniquely marked by individual ligands (Figure 6B-C; e.g. ChC: PTHLH, PV: Tac1, Adm, nNOS/SST: Ptn, Rln1, CR/SST: Nppc, VIP/CCR: Edn3, Pnoc). These results indicate that, beyond their morphological and physiological differences, GTPs are different neuroendocrine cells that produce distinct chemical outputs and elicit distinct physiological effects. Consistent with the demand for processing and packaging diverse neuropeptides, the granin gene family, which regulates pre-prohormone cleavage and biogenesis of DCVs (Bartolomucci et al., 2011), also shows differential expression among GTPs (AUROC=0.81 Table S4).

### Ptn in long projection GABA neurons may recruit oligodendrocytes for axon myelination

Although the function of most neuropeptides in GTPs are unknown, current knowledge on Pleiotropin (PTN) (Papadimitriou et al., 2016) enabled us to predict and then validate an unexpected cell phenotype in SST/nNOS long projection cells. PTN promotes axon myelination by activating the differentiation of oligodendrocyte precursors (Kuboyama et al., 2015). The unique expression of PTN in SST;NOS1 cells suggests that their axons might be myelinated. It is possible that PTN may promote LPC axon myelination during postnatal development and contribute to myelin maintenance in mature LPCs

To test this prediction, we examined the expression of CASPR (Gordon et al., 2014), a key component of the node of Ranvier, along LPC axons. Indeed, CASPR consistently co-aligned in a paranodal pattern along LPC axons (Figure 6E; Figure S6), demonstrating that these axons are indeed myelinated. This finding is surprising as cortical GABAergic interneurons are thought to elaborate unmyelinated axons that enable extensive branching and innervation of local target cells. But the unique feature and property of LPCs suggest that myelination of their long projecting axons, many extend through the white matter, may enhance their conduction speed to regulate global cortical networks (Kilduff et al., 2011; Tamamaki and Tomioka, 2010).

### Vesicular zinc transporter in SST/CR cells suggests a GABAergic synaptic source of zinc signaling

In addition to amino acid-based transmitters and modulators, the divalent cation zinc acts as a bona fide neuromodulator that exerts potent and pleiotropic impacts on neuronal signaling (Marger et al., 2014). Zinc is enriched in mammalian cerebral hemisphere, where the vesicular transporter ZnT3 in a subset of glutamatergic neurons loads synaptic vesicles for co-release with glutamate. Synaptic release of zinc modulates multiple ion channels, especially certain types of extra-synaptic NMDA receptors (Marger et al., 2014). In particular, activity-dependent increase of Zinc at synapses inhibits GluN2A-containing NMDARs at nano-molar potency, which impacts glutamatergic transmission, plasticity and circuit operation (Vergnano et al., 2014) (Romero-Hernandez, Furukawa 2016). It is unknown whether non-glutamatergic neurons mediate synaptic zinc signaling.

Surprisingly, we discovered that ZnT3 is highly and specifically expressed in SST/CR cells (Figure 6F). In addition, Zip1 and Zip7a transporters that mediate zinc uptake to the cytosol are also expressed in these cells. Therefore, SST/CR cells are equipped to accumulate cytosolic zinc for synaptic vesicle loading. These results suggest that SST/CR cells may co-release zinc and GABA. Most SST/CR cells are Martinotti cells (He et al., 2016) that target the distal dendrites and spines of pyramidal neurons with abundant GluN2A-NMDARs (Silberberg and Markram, 2007). Our results suggest that Martinotti cells might exert their powerful dendritic inhibition through two parallel mechanisms: synaptic activation of GABA_A_Rs with GABA and extra-synaptic inhibition of NMDARs with zinc. As Martinotti cells broadly innervate other types of GABAergic neurons (Jiang et al., 2015), similar mechanisms might mediate their non-selective and potent inhibition of GABAergic populations.

### Synaptotagmin members correlate with vesicle neurochemical contents

The current framework of the molecular machinery of vesicular release consists of SNARE complexes that form the core fusion pore, the Ca^2+^ sensing and regulatory components, the active zone that organizes the release site, and recycling of synaptic vesicle pools (Sudhof, 2013). The molecular components of vesicle fusion machinery are encoded by several multi-gene families, but it is not well understood how different members of these gene families shapes the release properties for transmitters and neuropeptides (Martin, 2003; Moghadam and Jackson, 2013; Sudhof, 2002).

Our transcriptome analyses reveal a comprehensive picture of the molecular profiles of vesicle release machinery in GTPs (Figure 6G-I). Several core components of the fusion complex and active zone are broadly expressed, including Syntaxins (AUROC=0.5), SNAP complex (AUROC=0.614), RIMs and RIM binding proteins (AUROC=0.57) (Table S5). Yet even among these core components, VAMP (synaptobrevins) and SNAP members are significantly enriched in specific GTPs (Figure 6G-H). More prominent patterns relates to the vesicular Ca^2+^ sensor synaptotagmins (Syt): 14 of the 17 Syts are differentially expressed among GTPs (Figure 6G-I; AUROC=0.78); individual neurons expresses 6-9 Syts (>30 uTPM; data not shown). In particular, VIP/CCK cells specifically express Syt10 that mediates the release of Igf1 (Cao et al., 2011), which is also highly enriched in the same cells (Figure 6D, G). These results suggest that in individual cell types and individual neurons, different Syts may act as Ca^2+^ sensors with different sensitivity to spatiotemporal Ca^2+^ signals that trigger particular types of fusion reactions, thereby contributing to the specificity and properties in parallel exocytosis pathways.

### Molecular signatures of vesicular release styles

In addition to chemical contents, the styles of transmitter release exert profoundly impacts on postsynaptic cells. The release sites of GABA and neuropeptides range from synaptic, axonal to somato-dendritic; the temporal characteristics range from fast, precise and synchronous (Hu and Jonas, 2014) to slow, sustained and asynchronous (Jonas and Hefft, 2010); and the short term dynamics following action potential trains range from facilitating to depressing (Markram et al., 2015). These different release styles produce distinct spatiotemporal patterns of receptor activation and postsynaptic cell firing that impact circuit level computation (Markram et al., 2015), but the underlying molecular mechanisms are not well understood.

Although it is often difficult to directly correlate molecular profiles of vesicle machinery to transmitter release properties, the extensive knowledge on GABA release styles of PV and CCK basket cells (Armstrong and Soltesz, 2012) provides a unique opportunity. Whereas PV cell-mediated GABA release is fast, precise and synchronous (Hu and Jonas, 2014), that of CCK cells is slow, sustained and asynchronous (Jonas and Hefft, 2010). A comparison of their transcription profiles begins to depict the molecular distinction of fast vs slow release machines, which manifests even at the level of core fusion complex. We found that VAMP1, SNAP25, Nsf and Rab3a are highly enriched in PV compared to VIP/CCK cells (Figure 6G-H); co-expression of all three components in CHC and PV cells supports their fast spiking and release properties (Parpura and Mohideen, 2008). Furthermore, among the 4 Ca^2+^-binding Syts localized to SVs (Syt1, 2, 9, 12), both PV and VIP/CCK cells (and other GTPs) express Syt1, but only PV cells express Syt2, which exhibits the fastest onset and decline in release. In addition, PV cell express highest level of Cplx1 but lowest level of Cplx2, while VIP/CCK cell profile is the exact opposite (Figure 6I-J). This Cplx profile is highly congruent with the Syt profile, as Cplx1 is implicated in fast and synchronous release and in clamping spontaneous release (Yang et al., 2013). Together, these results suggest that fast-synchronous release machine is built with high levels of VAMP1, SNAP25, NSF, Syt2, Cplx1. In contrast, slow-asynchronous release machine is built with low levels of these components and high levels of Cplx2. Together with expression of matching properties of Ca^2+^ channels and CaBPs, these results suggest that different cell types are endowed with distinct vesicular neurochemical profiles and their release styles, implemented by coordinated differential expression of gene families that encode components of the vesicle fusion machinery.

In addition to PV and CCK basket cells, it is notable that nNOS/SST long projection neurons express over 11 peptides (Figure 6B-D) and Syt4, 5, 6 are highly or uniquely enriched in these cells (Figure 6G). It is possible that these uncharacterized Syts might mediate the synaptic release of peptide-containing DVCs or their endocrine/paracrine release along the axon-dendritic membrane.

### Transcription factor profiles register the developmental history and contribute to the maintenance of GTP phenotypes

A recurring theme in our transcriptome analysis is the highly correlated and congruent differential expression among GTPs of select gene ensembles from gene families within and across the five functional categories (Figure 2D-E), suggesting orchestration by gene regulatory networks. Approximately 10% of the protein coding genes in the mouse genome are devoted to ~1,500 transcription factors (TFs). The combinatorial actions of TFs through their myriad cognate cis-regulatory elements provide enormous capacity for controlling the spatial and temporal transcription pattern of thousands of genes involved in the specification as well as the maintenance of cell identity (Davidson, 2010; Deneris and Hobert, 2014; Hobert, 2011), but the mechanisms that implement such transcriptional control are far from understood. As a prerequisite, comprehensive TF expression profiles in neuron types have not been described.

Here we report quantitative expression profiles of TFs in GTPs and explore their implications in the maintenance of cell phenotypes and (Figure 7). We found that each GTP on average expresses ~350-400 TFs and over 300 TFs are expressed in an individual cell (at >30uTPM in >75% of cells in each GTP; Figure 7C). Among ~34 TF classes, basic-helix-loop-helix (bHLH) proteins, nuclear hormone receptors, POU-homeoboxes, and kruppel-like transcription factors are most differentially expressed among GTPs (Table S5). A comparison between MGE vs CGE GABA neurons revealed that ~65 TFs are common between the two populations with ~90 TFs enriched in MGE and ~110 TFs enriched in CGE populations (at >30uTPM in >75% of cells in each GTP). Among the 4 MGE-derived and 2 CGE-derived GTPs, each expresses between 150-220 TFs (>30uTPM in >75% of GTP population, Figure S7) and multiple TFs individually marks each GTP (>2 folds enriched, at >30uTPM) (Figure 7H).

**Figure 7:**
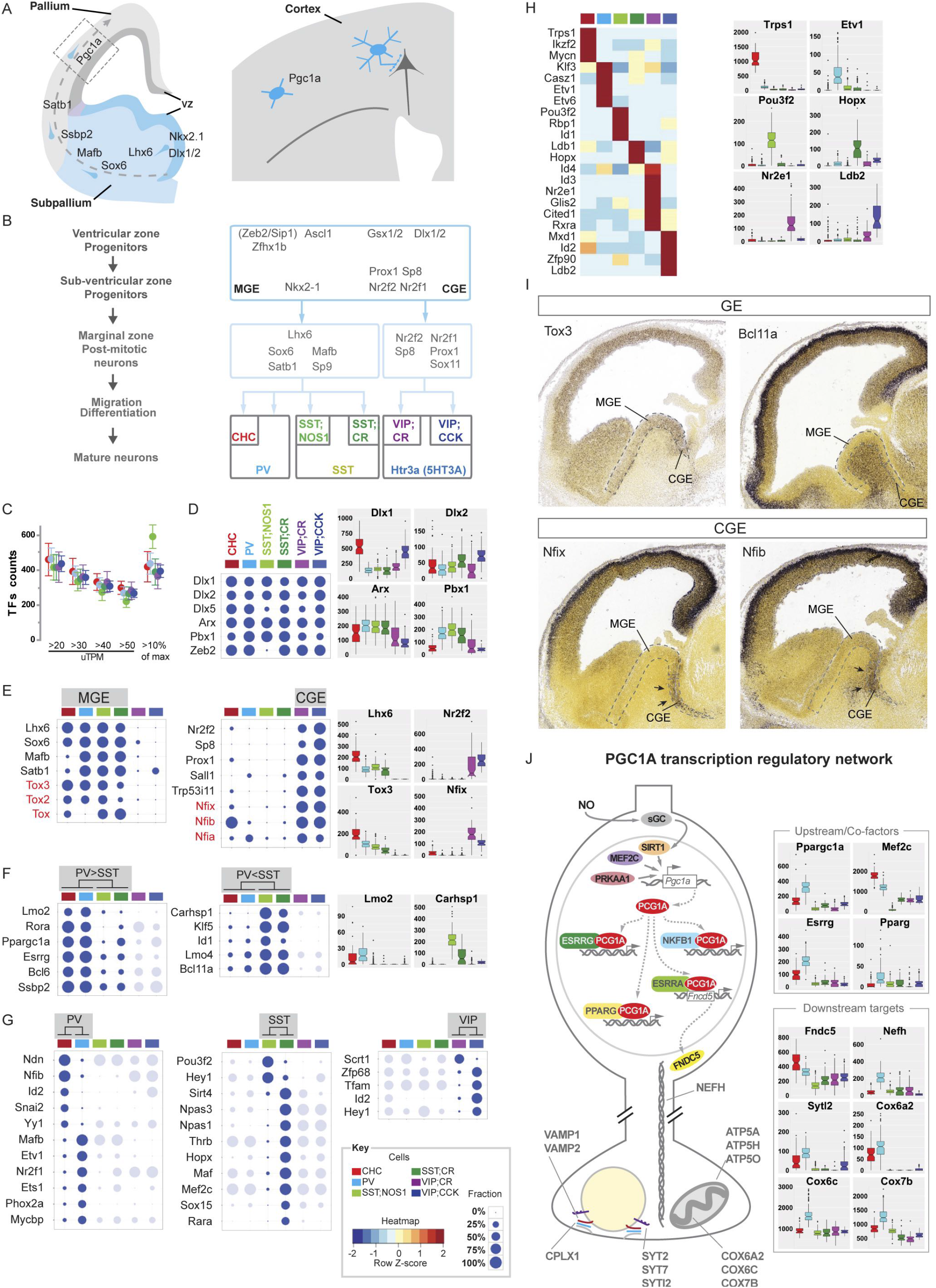
Transcription factor profiles register the developmental history and contribute to the maintenance of GTP phenotypes. **(A)** Schematic representation of the developmental trajectory of cortical GABAergic neuron, using PV cells as an example. TFs reported to express at different stages of PV cell development are shown. **(B)** Schematic summary of MGE and CGE transcription cascades that regulate the development of different clades of cortical GABAergic neurons including GTPs **(C)** Each GTP is estimated to express ~400 TFs based on sliding thresholds mRNA counts (2050 uTPMs) or dynamic thresholds (10% of maximum expression value). **(D-G)** Bubble-plot showing fraction of cells expressing a given TF at 10% of maximum expression value. Accompanying boxplots show expression levels of selected TFs, y-axis in uTPM. **(D)** Major TFs in subpallium progenitors and GABA neuron precursor maintain their expression in GTPs in adult cortex. **(E)** TFs expressed in early postmitotic MGE- and CGE-derived neuron maintain expression within same clade of GTPs. Embryonic expression of Tox and Nfi family TFs were deduced from our transcriptome analysis of GTPs and confirmed in (I). **(F)** Among MGE-derived GTPs, subsets of TFs are preferentially expressed in PV (PV>SST) or SST (PV<SST) subpopulations; CGE groups are not compared and shown in light shade. Representative boxplot shown to the right. **(G)** Within the PV, SST and VIP group, subsets of TFs are enriched in one or the other GTP; GTPs that are not compared are shown in light shade. **(H)** Heatmap showing differential expression of TFs largely exclusive to each GTP; boxplots show several examples of ON/OFF TF expression exclusive in a GTP at indicated levels (uTPM). **(I)** Retrospective screen of Allen Developmental Mouse Brain in-situ database reveals that several TFs that express in MGE- or CGE-derived GTPs identified by transcriptome analysis indeed begin their expression in the corresponding embryonic germinal zone. **(J)** Schematic of the Ppargc1α (PGC1α) transcription regulatory network which is highly enhanced in PV cells. Multiple reported PGC1α upstream TFs activators, cofactors and large fraction of (>75%) of reported downstream effectors are enriched over 1.5 folds in PV cells (p<5.0^-07^). See Figure S7 for a detailed comparison of known PGC1α targets and their differential expression in PV cells. Boxplots show different expression levels of select sets of PGC1α co-factors and targets and putative targets among GTPs.

### GABAergic neurons retain a transcription resume that registers their developmental history

The molecular mechanisms that maintain GABAergic cell phenotypes and identities are poorly understood. Studies in invertebrates and other mammalian brain regions suggest that neuronal identity is maintained in postmitotic cells by sustained expression of the same set of transcription factors that initiate terminal differentiation during neural development (Dalla Torre di Sanguinetto et al., 2008; Deneris and Hobert, 2014). Our experimental design allowed us to examine this hypothesis by comparing TF profiles in GTPs with those in their embryonic precursors during development.

The embryonic subpallium contains a developmental plan embedded in progenitors along ganglionic eminence whereby transcription cascades orchestrate the specification and differentiation of major clades (i.e. MGE, CGE) of cortical GABAergic neurons (Kepecs and Fishell, 2014; Nord et al., 2015) (Figure 7A-B). A unique feature of our experimental design is that the 6 GTPs are embedded in 3 non-overlapping populations (PV, SST, VIP) from 2 separate developmental origins (MGE vs CGE) (Figure 7B). This data structure provides a developmental context for establishing the link between TF profiles in mature neurons to those in their embryonic precursors with cell type and single cell resolution. We found that almost all the well-studied TFs in embryonic precursors maintain their expression within the same clade of mature GTPs (Figure 7D, E): whereas Lhx6 (Du et al., 2008; Zhao et al., 2008), Sox6 (Azim et al., 2009; Batista-Brito et al., 2009), Mafb (McKinsey et al., 2013), Satb1 (Close et al., 2012; Denaxa et al., 2012) are expressed in PV and SST populations (i.e. the MGE clade), Coup-TF2 (Lodato et al., 2011), Sp8 (Ma et al., 2012), Prox1 (Miyoshi et al., 2015), Npas1, Npas3; Stanco et al., 2014) are expressed in VIP populations (i.e. the CGE clade). Furthermore, we discovered multiple additional TFs with similar patterns: whereas Tox family members (Tox, Tox2, Tox3) (Artegiani et al., 2015; Sahu et al., 2016) are restricted to the MGE clade, Nfi family members (Nfia, Nfib, Nfix) (Mason et al., 2009; Piper et al., 2014), Sall1 and Trp53i11 are restricted to the CGE clade. Importantly, by “reverse tracking” of their developmental history through screening the Allen Developmental Mouse Brain Atlas, we found that each of these TFs is indeed expressed in the embryonic MGE or CGE, consistent with their clade relationship (Figure 7I).

Furthermore, by hierarchical and pair-wise comparison, we defined multiple sets of TFs that distinguish PV vs SST population (Figure 7F), and ChC vs PV, SST/nNOS vs SST/CR, and VIP/CR vs VIP/CCK cells (Figure 7G). Again, we found evidence for developmental continuity of TF expression from embryonic precursors to mature neurons (also see Supplemental Text 4 and Figure S7B), with PV cells as the most notable case. Multiple PV-enriched TFs initiate expression at different stages of development, from Lhx6 and its downstream cascade (Figure 7A, D-E) to Ssbp2, Nfib and Etv1 (Figure 7E-G; (Batista-Brito et al., 2008). In addition, the Ppargc1α (peroxisome proliferator-activated receptor γ coactivator 1α) expression initiates in the first few postnatal days and maintains expression in mature PV cells (Cowell et al., 2007; Lucas et al., 2010). Although the comprehensive developmental profile of TF expression in PV cells remains to be determined, current evidence indicates that PV cells maintain sequentially acquired transcriptional programs, from early genetically determined TFs to late activity-regulated TFs (Figure 7A).

Together, these results suggest that mature GTPs register the developmental history of their transcription program, i.e. a “transcription resume”, and the sustained expression of sequentially accumulated transcription programs might contribute to the maintenance of corresponding cell phenotypes and properties. Consistent with this notion, conditional deletions of several of these TFs in adult cortex result in the loss of markers and physiological properties characteristic to their identity (Batista-Brito et al., 2009; Close et al., 2012; Miyazaki et al., 2012; Touzot et al., 2016), suggesting that they are required in mature neurons to maintain specific aspects of cell phenotypes and identity.

### The PGC1α transcription program coordinates the release and metabolic properties of PV cells

Consistent with the fast input-output transformation of PV basket cells, our analysis revealed highly congruent expression of fast-acting molecular components at every step of electrical signaling (e.g. Figure 4C-I, Figure 6G-K). In addition, the operation of PV cells requires specialized energy metabolism and mitochondria features (Lucas et al., 2010). It is unknown whether the molecular components mediating these distinct yet functionally congruent properties are cobbled together in a piece meal fashion or are orchestrated by transcription programs.

The transcription coactivator PGC1α was discovered in peripheral tissues and cooperates with multiple transcription factors to regulate mitochondria biogenesis and energy metabolism in response to fasting and exercise (Lin et al., 2005). In the brain, it is restricted to subset of GABAergic neurons, especially PV cells in the neocortex (Cowell et al., 2007). During brain development, PGC1α expression increases dramatically in the first and second postnatal week, corresponding precisely with the substantial morpho-physiological maturation of PV cells during this period, and remains in PV cells thereafter (Lucas et al., 2010). Microarray studies in cell cultures and gene deletion in mice revealed that PGC1α likely directly regulates the expression of PV and genes involved in mitochondria function and transmitter release (Lucas et al., 2014). However, the transcription co-factors and comprehensive transcription target genes of PGC1α in PV cells in vivo are unknown. Our transcriptome analysis revealed that, in addition to PGC1α, several of its co-factors (e.g. Rora, Esrrg, Pparg; Figure 7J) are significantly enriched in PV cells, suggesting a specific enhancement of the PGC1α transcription program. Furthermore, we confirmed that all the potential targets from previous studies (Lucas et al., 2014) are substantially enriched in PV cells (PV, Syt2, Nefh, Cplx1, Atp50, Atp5a1). In addition, we found an extended set of PV enriched mRNAs associated with metabolic and mitochondria pathways (e.g. Fndc5, Cox6a2, Cox6c, 7b; Figure 7J). Although a proof of transcriptional targets would require evidence of their alteration in PGC1α deficient cells and, ultimately, of direct binding to cis-regulatory elements, our results suggest that PGC1α may coordinate a transcription module or gene battery that shapes the release and metabolic properties in PV cells.

## Molecular portraits of GABAergic cell types

We have discovered highly correlated and congruent gene expression across multiple gene families and functional categories in each GTP. While coordinated expression within a functional category (e.g. voltage-gated ion channels) may shape a specific cell property (e.g. fast spiking), those across functional categories may shape a set of congruent cell properties that together characterize cell identity (e.g. fast input-output transformation in PV cells). Here we highlight the coherent and synergistic nature of these expression profiles that may to GTP phenotypes (Table 1).

### PV basket cell (PVC)

PVCs innervate the perisomatic region, mediate fast input-output transformation, and contribute to feedforward, feedback inhibition and fast network oscillations (Hu et al., 2014). Their weakly excitable dendrites feature fast glutamate receptors, high levels of Kv3 channels, and very low level Navs; together these channel composition and distribution patterns allow high activation threshold, fast activation and fast de-activation. On the other hand, their axons feature several Na channel members (e.g. Nav1.1, Nav1.6) at exceedingly high levels and multiple types of K channels (e.g. Kv1); these ensure fast action potential generation and reliable and fast propagation. Furthermore, their axon terminal features fast gating P/Q type Ca^2+^ channels tightly coupled to release machinery, the fastest Ca^2+^-sensing synaptotagmin (Syt2), and high concentration of calcium binding proteins (e.g. PV); together these ensure minimum synaptic delay for fast GABA release.

Our transcriptome analysis confirmed all previously reported PVC markers (Table 1 in (Hu et al., 2014) and further allow quantitative comparison with other cell types with different physiological properties (e.g. CCK basket cells) to infer the significance of differential gene expression (e.g. Figure 6K). More importantly, we discovered a much larger set of PVC-enriched mRNAs that may contribute to their known properties and predict novel properties. With regard to voltage-gated ion channels, we found enrichment of Nav1.7 and a large set of regulatory subunits of Nav (Scn1b, Scn4b), Kv (Kcnab1, Kcnab3) and Cav (Cacna2d2) that likely co-assemble with corresponding core subunits to shape native channel properties. Regarding input, GluR core and auxiliary subunit profile (e.g. Cacng2, Figure 4E) suggests specialized native receptors likely endowed with specific trafficking and biophysical properties. Furthermore, GABA_A_R profile predicts the highest level and variety of receptor subtypes among GTPs (Figure 4H-I). Regarding output, the striking co-enrichment of Vamp1, Syt2, Syt7, Cplx1, Snap25, NSF depicts increasing details of a fast-synchronous release machine (Figure 6G-K), and expression of novel neuropeptides (e.g. Adm, Rspn) suggests release of additional ligands (Figure 6B-D). Furthermore, novel signaling proteins (e.g. RGS4, Adcy8, Rasl11b, Arhgef10l) predicts specialized intracellular signaling properties (Figure 5B-D). Although expression of cell adhesion molecules are difficult to interpret, the specific enrichment of Slit and Unc5 members (Figure S3A) suggest their roles in axon branching, cell recognition and wiring specificity. Finally, our analysis of PGC1α transcription program revealed putative co-transcription regulators (Esrrg, Mef2c; Figure 7J) and more comprehensive transcription targets (e.g. Fndc5, Cox6a2, Cox6c) that likely support the metabolic demand and enhanced mitochondria function.

Together, these results begin to depict a substantially high resolution molecular portrait of PVCs that contribute to fast signaling at every step: from fast excitation, fast-narrow spiking, to reliable spike propagation, fast GABA release and metabolic support. Such congruent expression of a large set of gene ensembles across multiple functional categories cannot be explained without invoking a master transcription program.

### CCK basket cell (CCKC)

CCKCs are enriched in the VIP/CCK subpopulation. Although these cells also innervate the perisomatic region of pyramidal neurons, they show different, often opposite, physiological properties compared to PVCs (Armstrong and Soltesz, 2012). CCKCs receive less glutamatergic inputs but rich neuromodulatory inputs that convey mood and brain states. Accordingly, here we show that CCKCs express much less GluRs but a large set of modulatory receptors. In addition to the reported endocannabinoid (CB1R), serotonin (Htr2c, Htr3a) and acetylcholine (Chrm3) receptors, they further express neuropeptide (Vipr1, Npy1r) and norepinephrine (Adra1b) receptors (Figure 4). Contrary to PVCs, CCKCs mediate slow, asynchronous and sustain GABA release; these properties likely derive from the profile of their release machinery, which features low level of Vamp1, Syt2, Syt7, Cplx1, Snap25, NSF but high levels of Cplx2 (Figure 6G-K). Furthermore, CCKCs express a large set of neuropeptides and hormones (>12) and at least one of these, Igf10, is likely mediated by a specialized synaptotagmin (Syt10) (Figure 6B, D, G). Together, these results depict a molecular profile highly distinct from PVCs: CCKCs likely integrate information from multiple modulatory systems over longer time windows, exert more sustained inhibition to the perisomatic region, and release multiple neuropeptides and hormones through specialized release machineries.

### Chandelier cell (ChC)

Although considered similar to PVC in several aspects, such as fast spiking and controlling pyramidal neuron output at the perisomatic region, ChCs are unique in their exquisite specificity to pyramidal neuron and to AIS, suggesting highly specialized function. Superimposed upon the notable similarity in their transcriptome profiles, we found significant differences across multiple gene categories between ChCs and PVCs. ChCs have: 1) different set of adhesion molecules (e.g. Unc5b) and sulfotransferases (e.g. Hstst4) that might contribute to different connectivity and extracellular matrix; 2) lower levels of Nav1.6, Kv1 and Kv3.2, consistent with their “not-as-fast” spiking property; 3) very low levels of α1, α4, α5 GABA_A_Rs, suggesting that they receive weaker inhibitory inputs, especially from PVCs which mainly use α1-containing GABA_A_Rs; 4) enhanced NO and cGMP signaling (Gucy1b3, Prkg1, Pde11a, Pde5a); 5) distinct neuropeptide profiles; 6) distinct release machineries characterized by contrasting pattern of Vamp1, Vamp2, Syt1, Syt2, Syt7, Cplx1 – suggesting that ChC terminals do not mediate fast GABA release. Together, these results suggest that ChCs are distinct from PVCs in anatomic and physiological properties and represent a unique microcircuit module that exerts specific control of pyramidal neuron firing (Lu et al 2016 in revision).

### Martinotti cell (MNC)

MNCs are enriched in the SST/CR subpopulation. They innervate the distal dendrites of pyramidal neurons and many other GABAergic neurons (Jiang et al., 2015; Silberberg and Markram, 2007). They exhibit high spontaneous activity in vivo during resting state (Gentet et al., 2012) and provide powerful inhibition to broad targets, with an exception of other MNCs (Jiang et al., 2015). With sparsely spiny dendrites, their hippocampal homologue (O-LM cells) manifests AMPAR-mediated plasticity somewhat similar to that of glutamatergic pyramidal neurons (Oren et al., 2009). Our transcriptome analysis begins to suggest some underlying molecular mechanisms. Prominent transcription features of MNCs include: 1) highest level and largest variety of AMPARs and auxiliary subunits, 2) highest level and variety of NMDARs, 3) least variety and lowest levels of GABA_A_Rs. Interestingly, Grin3a is in fact a glycine receptor (Cummings and Popescu, 2016; Perez-Otano et al., 2016) that mediates depolarization and may contribute to high excitability (e.g. “low-threshold spiking”; (Amitai et al., 2002) and spontaneous tonic firing (Gentet et al., 2012). High levels of Gria1a, Grin1, Grin2a and low levels of GABA_A_Rs may contribute to enhanced glutamatergic excitability and plasticity (Oren et al., 2009). The most surprising finding is the specific expression of vesicular zinc transporter (Figure 6F), suggesting co-release of GABA and zinc that mediate two parallel inhibitory pathways: synaptic activation of GABA_A_Rs with GABA and extra-synaptic inhibition of NMDARs with zinc. Together, our results suggest mechanisms for a cell type with high level tonic firing and excitability, enhanced glutamatergic plasticity for learning, low level inhibition in part mediated by disinhibition, and co-release of GABA and zinc that might represent a novel mechanism to coordinate GABAergic and glutamatergic synaptic transmission and plasticity.

### Interneuron selective cells (ISC)

The VIP/CR population contains ISCs that mainly innervate SST and PV interneurons but avoid pyramidal neurons. ISCs can be recruited by several modulatory transmitters that signal salient sensory or internal stimuli to mediate rapid disinhibition of subsets of pyramidal neurons (Pfeffer et al., 2013; Pi et al., 2013). The prominent expression of multiple neuromodulatory receptors (Htr2c, Htr3a, Chrm3, Chrma4, Adra1b) is consistent with previous findings and predict novel input sources. Their release machine profile is similar to CCKCs, suggesting asynchronous, sustained GABA release that may mediate longer lasting disinhibition.

### Long projection cell (LPC)

Among the 6 GTPs, type I NOS long projection neuron is the most unusual and least understood. These cells appear to increase activity following sleep deprivation-induced recovery sleep and have been suggested to regulate global cortical states (Kilduff et al., 2011), but their cellular properties and physiological function are largely unknown. Our transcriptome analyses reveal and predict numerous physiological and anatomical features that can be validated. In terms of input, these neurons show the lowest level of iGluR core subunits (Figure 4C-E), suggesting weak glutamatergic excitation. They express highly unusual form of γ1 containing, likely non-synaptic, GABA_A_Rs (Figure 4H-I), suggesting slow GABAergic inhibitory inputs. Thus LPCs appear less engaged by fast glutamate and GABA transmission systems. Instead, they express a large set of neuromodulatory GPCRs, including acetylcholine, 5-HT, oxytocin, hypocretin (sleep peptide), and the unusual extra-retinal photoreceptor opn3 (Figure 4K-M). These results indicate that LPCs are strongly influenced by brain state modulatory inputs. In terms of output, in addition to GABA and NO, they express the largest number (~14) neuropeptide mRNAs (Figure 6B-C). Accordingly, they express multiple Ca^2+^-insensitive synaptotagmins (syt4, 5, 6), which may contribute to non-synaptic peptide release through dense core vesicles (Figure 6G). In terms of signaling, they feature the entire NO signaling pathway, from substrate accumulation and NO synthesis to 2^nd^ messenger cascade and target effectors (Figure 5E-F). A major surprise was our discovery, based on prediction from the specific expression of Ptn, that LPC axons are myelinated (Figure 6E). In contrast to most GABAergic interneurons with local axon arbors that are mostly unmyelinated, the myelination of LPC axons may increase the conduction velocity to their distant terminals. Together, these depict a large neurosecretory cell type that can be recruited during various behavioral states to modulate a wide range of neuronal and nonneuronal targets in global cortical networks. The highly coherent co-expression of multiple sets of functionally congruent gene ensembles cannot be explained without invoking an underlying transcription program.

## DISCUSSION

Understanding the nature of neuronal identity is prerequisite to deciphering brain cell diversity and has remained a fundamental unsolved issue in neuroscience since the discovery of the Neuron Doctrine. Despite substantial progress in characterizing cell phenotypes along multiple axis and in cell type classification in several invertebrate species (e.g. worms, flies) and brain regions (e.g. the retina), the molecular biological basis of neuronal identity has remained obscure and a mechanistic framework has been elusive. Here, through single cell transcriptome analysis of anatomically and physiologically characterized ground truth cortical GABAergic neurons, we have discovered the transcriptional architecture of cardinal neuron types. This genetic architecture mainly encodes synaptic connectivity and input-output properties and comprises primarily 6 functional categories of ~40 gene families that include cell adhesion molecules, transmitter-modulator receptors, ion channels, signaling proteins, neuropeptides and vesicular release components, and transcription factors. Combinatorial and congruent expression of select members across these families is likely orchestrated by cell specific transcription programs, which shape a multi-layered molecular scaffold along cell membrane and may customize the patterns and properties of synaptic communication. In essence, our results suggest that the core identities of neuron types are encoded in key transcription programs that organize and constrain their basic patterns of physiological connectivity. As connectivity is the primary purpose of morphology and physiological connectivity is the basis of circuit operation, this overarching and mechanistic definition integrates the molecular, anatomical, physiological and functional descriptions of neuron types. Characterization of key transcription profiles of increasing number of ground truth types will provide reference axes for large scale transcriptome-based discovery of neuron types.

### Cortical GABAergic neurons are well suited for transcriptomic analysis of cell phenotypes

Compared with anatomical and physiological descriptions of cell properties, scRNAseq provides high dimensional, quantitative and high-throughput measurements of mRNA profiles, but transcriptome dataset needs to be validated by, harmonize with and explain orthogonal cell phenotypes that more directly contribute to circuit function. In most brain regions, knowledge of cell phenotypes are often grossly incomplete and sketchy, thus transcriptome data may rapidly accumulate without the complementary support of and validation by neurobiological information. In this context, cortical GABAergic neurons are particularly well suited for exploring the molecular genetic basis of cell type identity through transcriptomics. Over 3 decades of anatomical, physiological, molecular and developmental studies have accumulated substantial ground truth information on a large set of GABAergic populations and in some cases bona fide cell types (Hu et al., 2014; Huang, 2014; Jiang et al., 2015; Kepecs and Fishell, 2014; Markram et al., 2015). Importantly, the massive use of subcellular patch-clamp techniques and multiple cell recording provide quantitative physiological measurements of intrinsic, synaptic and release properties that more directly relate to gene expression and molecular mechanisms (Hu et al., 2014; Jiang et al., 2015; Markram et al., 2015). This information is further enriched by leveraging on deeper understanding of homologous cell types in the hippocampus (Somogyi et al., 2014). Therefore, cortical GABAergic neurons present a rare opportunity to systematically validate and harmonize gene expression profiles with specific morphological, biophysical, connectivity phenotypes. These ground truth information are essential for recognizing technical noise, statistical pitfalls and guiding our computational analysis. Indeed, many of our results are readily validated by the literature, provide deeper explanation of known cell phenotypes, and predict novel properties that can be examined with cell targeting tools (He et al., 2016).

### Computation genomics screen of gene ensembles that contribute to cell phenotypes

Our study differs from and complements several recent single cell transcriptome analyses (Shekhar et al., 2016; Tasic et al., 2016; Zeisel et al., 2015). Whereas previous studies aimed to classify cell diversity using statistical clustering of transcriptomes from unbiased cell populations (Zeisel et al., 2015) or relatively broad populations (Tasic et al., 2016), our goal was to discover the molecular basis of cell identity by analyzing phenotype-characterized subpopulations and bona fide cell types. In terms of methods, we combined 2-round of linear amplification of mRNA with unique molecular identifiers, which achieved more comprehensive coverage (>8000 genes) and more quantitative measurements with a higher dynamic range (Figure S1) that facilitate subsequent computational analysis. Previous analyses of differential gene expression often identify sets of molecular markers for “transcriptomic types”; although useful, these markers often appear piece meal, serendipitous, and do not readily inform cell phenotypes. On the other hand, we focused on analyzing differential gene ensemble profiles (i.e. gene families) encoding proteins that constitute cellular modules (Hartwell et al., 1999) (i.e. macromolecular machines, signaling complexes and pathways), which more readily relate to, explain and predict cell phenotypes. Most importantly, using ground truth population as an *assay*, we designed a supervised machine learning-based computational genomics strategy to systematically *screen* all gene families and rank their abilities to discriminate GTPs. This method enabled us to discover a rather small set of functionally related gene families, which constitute a coherent transcriptional architecture that organizes the physiological connectivity patterns that characterize GTPs. The MetaNeighbor (Crow et. al. 2017) method can be applied to other gene ensembles such as GO terms, custom-annotated gene groups for signaling complexes and pathways to extract knowledge and insight from transcriptomic dataset.

### Transcriptional architecture of physiological connectivity defines neuronal cell types

An enduring challenge in classifying neurons in mammalian brain is that most if not all methods only provide sparse, partial and often less than quantitative measurements of 1-2 aspects (e.g. morphology, electrophysiology) of the inherently multi-faceted neuronal phenotypes. Thus the discordance between “cell types” defined by sparse and incomplete datasets on different phenotypes (DeFelipe et al., 2013; Petilla Interneuron Nomenclature et al., 2008) should perhaps not be surprising. Combining deep and quantitative transcriptomic with a computational screen that links gene ensembles to phenotype-characterized ground truth cell populations, our conclusion that transcriptional architecture of physiological connectivity defines neuron types can serve as an integrated and overarching scheme for neuron classification for several reasons. First, although morphology is the most common and intuitive description of neurons, it is increasingly recognized that morphology reflects and serves the more fundamental purpose of connectivity (Seung and Sumbul, 2014). Indeed, morphological types can be reliably identified from dense connectomes, when available, by computational algorithms (Jonas and Kording, 2015). Furthermore, the variability of morphology likely belies the co-variation of pre- and post-synaptic neurites that preserve the same connectivity pattern (Sumbul et al., 2014). Therefore connectivity is the primary purpose of anatomy that integrates cell position and neurite morphology. Beyond anatomical connectivity, the functional purpose of a neuron type is to transform the physiological contents embedded in its synaptic inputs (e.g. excitatory vs inhibitory vs modulatory nature, strength, dynamics, spatiotemporal patterns) into appropriate outputs (Kepecs and Fishell, 2014; Sharpee, 2014), often characterized by cell intrinsic style of neurochemical release (transmitter-modulator contents, speed, dynamics, plasticity). Although extremely valuable, most electrophysiological measurements are made at cell soma regions, often in artificial conditions, that provide a rather limited window into the elaborate subcellular biophysical, cell biological and metabolic processes. Our analysis provides a comprehensive transcription overview of the synaptic, intrinsic and release machineries and reveals strikingly coherent molecular properties congruent with well characterized intrinsic, synaptic and release properties of GTPs (e.g. PVCs). They further predict multiple novel physiological features that can be experimentally verified. Thus transcription signatures of synaptic input-output machineries may begin to harmonize with and integrate the hitherto often limited, disparate and technically challenging electrophysiological measurements.

In considering circuit function, cell types defined by their synaptic connectivity and input-output styles represent distinct structural and physiological motifs, with characteristic and restricted set of dynamic properties that support and constrain their role in circuit operations (Huang, 2014; Kepecs and Fishell, 2014; Markram et al., 2015). Differential assembly and physiological recruitment of these motifs into larger networks engage their participation in systems level information processing that underlies various context-dependent behaviors. In this sense, the physiological connectivity definition of cell types may integrate multi-parametric morphological, physiological and functional descriptions.

With current technical limitations, neuron type definition by physiological connectivity will remain a working hypothesis to be tested by the accumulation of connectome dataset in vertebrate brains in addition to those in worms, flies and the retina (Chen et al., 2006; Helmstaedter et al., 2013; Takemura et al., 2013). Comprehensive measurements of input-output properties are even more challenging and likely require novel methods for large scale recording (Marder, 2012). On the other hand, the rapid progress of transcriptome analysis will likely reinforce our finding on the transcriptional architecture of cell types and provide deeper and increasingly mechanistic insights into cell identity from the perspective of developmental genetics. There has been much debate on the arbitrary vs biological nature of cell types and the operational vs mechanistic basis of cell classification especially for cortical interneurons (Battaglia et al., 2013; DeFelipe et al., 2013). Despite the inherent variability of cell phenotypes - some seemingly a continuum without apparent boundary (Battaglia et al., 2013) likely in part due to technical limitations for measuring core phenotypes (e.g. connectivity), it is notable that the primary features of bona fide cell types (e.g. chandelier cells) are nearly identical among individuals of the same species and are conserved across species (Woodruff and Yuste, 2008). This self-evident fact suggests that, despite the diversity and variability, cardinal neuron types are reliably generated through developmental genetic mechanisms that specify cell fate and constrain their differentiation potential toward their connectivity-based identity in neural circuits. Our discovery of the transcriptional architecture of GTPs is likely a manifestation of transcription programs shaped by developmental genetic mechanisms rooted in the genome.

### Transcription resume may reflect the developmental accumulation of gene regulatory programs that maintain cell phenotypes

The highly coherent co-expression of large sets of gene ensembles encoding functionally congruent protein complexes, signaling and metabolic pathways that synergistically shape characteristic properties of each GTP strongly suggest the orchestration by gene regulatory networks. Transcriptional control of “gene batteries” has been demonstrated in experimental systems where brute force genetic analysis can establish causal links between specific transcription factors and their target effectors (Hobert et al., 2010). Complementing and substantiating previous studies of the PCG1a transcription program in PVCs (Lucas et al., 2014), our transcriptome analyses suggest potential transcription co-regulators and additional putative target effectors (Figure 7J). Together these evidence suggest that, similar to the PGC1α program, many other transcription modules remain to be discovered that coordinate the expression of functional gene ensembles (e.g. NO and cGMP signaling pathway in SST/nNOS cells).

Furthermore, we found that numerous developmental transcription programs initiated at successive stages of postmitotic differentiation are maintained in the same clade of mature neurons (Figure 7A-G). Interestingly, conditional knockout of several such TFs result in the loss or alteration of interneuron identity consistent with their developmental expression and clade relationship (Close et al., 2012; Touzot et al., 2016). During the specification of diverse spinal motor neurons, a hierarchy of transcription programs progressively restrict cell fate and establish their competence in responding to subsequently initiated differentiation programs, including activity-dependent processes (Dalla Torre di Sanguinetto et al., 2008). It is possible that a similar developmental principle guide the postmitotic migration, differentiation and maturation of cortical GABAergic neurons. In addition, our results suggest that most developmental programs remain in mature neurons and constitute a transcriptional resume that may maintain cell phenotypes and identity.

Although our data are mostly correlative and causal links require experimental manipulations, the cumulative overwhelming evidence precludes piece meal, coincidental mechanisms of coexpression and leaves little doubt for orchestration of transcriptional programs that maintain cell phenotypes and identity. One of the greatest challenges in modern biology is to discover the transcriptional and epigenomic mechanisms that control gene expression programs underlying cell identity. Our study identifies a large set of co-regulated functional gene ensemble in multiple GTPs and provides opportunities to explore the underlying transcription factors and enhancers.

### Parallel GABAergic and peptidergic identities and their functional implications

Grounded on the concept that GABA is the major inhibitory neurotransmitter in vertebrate brains, investigations of cortical inhibitory neurons have centered around GABA-associated properties and function. Although neuropeptides are among the first set of serendipitous markers found in GABAergic neurons (Somogyi et al., 1984), their significance and implication have remained obscure. Our transcriptome analysis reveals that all GTPs are peptidergic and each has a distinct profile consisting of some 6-14 endogenous ligands (Figure 6B-C); indeed this gene family ranks at the very top in discriminating GTPs. Consistent with the fact that neuropeptides and hormones are differentially processed by proteolytic machinery and released from dense core vesicles with specialized fusion machinery, granin (implicated in neuropeptide processing and packaging (Bartolomucci et al., 2011)) and synaptotagmin (including Ca-independent variants implicated in hormone release) (Moghadam and Jackson, 2013) members are also differentially expressed among GTPs. The obligatory and extensive GABA-peptide co-expression indicates that cortical GABAergic neurons are inherently neuroendocrine cells. Furthermore, neuropeptide profiles and their packaging and release styles - key determinants of neurochemical output qualities, appear to constitute a peptide code of GABAergic subtype identity (e.g. chandelier cells, long projecting cells). These findings suggest that cortical GABAergic neurons are hybrid endocrine neurons with parallel identities represented by separate intracellular biogenic and release machines, intercellular signaling mechanisms, and physiological functions.

The pan-GABA identity is conferred by GABA synthesis (Gad1, Gad2), synaptic vesicle loading (vGAT) and release/recycling machineries. Diverse GABAergic neuron types manifest characteristic synapse specificity, input reception properties, intrinsic physiology and GABA release styles, all serving to craft specialized roles in fast, precise and dynamic inhibitory control of postsynaptic neurons and network activities through GABA receptors (Somogyi et al., 2014). On the hand, neuroendocrine identity is characterized by peptide synthesis, proteolytic processing and packaging through endoplasmic reticulum into dense core vesicles, and all-or-none type release machines (Moghadam and Jackson, 2013). Specific sets of peptides in GABAergic types are likely released from synaptic as well as non-synaptic sites following different firing patterns that trigger GABA transmission; they may act as separate streams of neuromodulators to transform neuronal intrinsic and synaptic properties through cognate GPCRs and reconfigure networks at a slower temporal window and broader spatial scale, as demonstrated in better understood systems (Marder, 2012).

Fast GABA-mediated inhibitory transmission and modulatory neuroendocrine signaling also have different evolutionary histories. Diverse neuropeptides are the predominant means of neurotransmission and modulation in invertebrate neural circuits (Bargmann, 2012)(Marder, 2012). Many of these peptide systems are conserved in vertebrate peripheral and central nervous systems including the mammalian forebrain. Indeed the genetic programs that configure neuropeptide biogenesis, release and signaling have evolved since early metazoans (Mirabeau and Joly, 2013). On the other hand, although GABA-mediated neurotransmission is found in worms, flies and other invertebrates, their genomes typically feature a gene pair encoding 2 GABA receptors (Tsang et al., 2007) that mostly form homo or hetero-meric cation channels instead of pentomeric anion channels (Beg and Jorgensen, 2003; Gisselmann et al., 2004). It is not until early vertebrates, when the emergence of 19 GABA_A_ receptor genes likely enabled the combinatorial assembly of highly diverse pentomeric GABA_A_ receptors that mediate fast chloride-based inhibitory transmission (Tsang et al., 2007). Notably, it is also in early vertebrates that the duplication and evolution of Dlx genes (Neidert et al., 2001) may have allowed their recruitment into ventral telencephalon progenitors of the ganglionic eminence and begin to drive forebrain GABAergic identity; this may further provide the basis for MGE and CGE genetic programs to specify forebrain GABAergic cell types (Nord et al., 2015). These considerations suggest at least two scenarios that cortical GABAergic endocrine cell types might arise: by building pan-GABA and GABA subtype identities upon a more ancient neuroendocrine identity template, or by initiating a novel forebrain GABAergic developmental program that incorporates available neuroendocrine genetic programs. One possibility is that GABAergic neuroendocrine identities are orchestrated by hierarchical transcriptional programs that specify and integrate GABAergic vs peptidergic features. For example, downstream to an overarching Dlx-based GABA identity, the PGC1α program in PVCs may coordinate fast GABA release and high metabolism while a separate yet coupled transcription program may regulate Tac1 and Adm biogenesis and release to endow a peptidergic profile in the same cell type. This hypothesis can be tested by identifying transcription factors that regulate and coordinate GABAergic and peptidergic features.

## Conclusions

We have discovered that the transcriptional architecture underlying synaptic connectivity and communication defines cardinal neuron types. Although transcription is influenced by cellular milieu including neural activity, core features of transcriptomes are outputs of cellular epigenomes customized primarily through developmental programming of the genome. Our finding thus suggests that the basic identities of cardinal neuron types are deeply encrypted in the genome. Characterization of key transcription profiles of increasing number of ground truth neuron types will provide valuable reference axes for large scale transcriptome-based discovery of cell types and facilitate identification of homologous cell types across species. As high resolution molecular portraits of neuron types predict connectivity and physiological properties with impact on circuit operations, transcriptome-derived insights on cell types will integrate molecular and systems level analysis of neural circuits. It is notable that ChCs and PVCs are implicated in the pathophysiology of schizophrenia (Gonzalez-Burgos et al., 2015), and many mental disorders likely result from altered neural connectivity linked to deficient cell type components. High resolution transcriptome analyses of neuron types help understand the molecular basis of cellular connectivity and physiology, facilitate linking altered gene expression to aberrant cellular and circuit properties that contribute to illness, and will provide new set of therapeutic targets to ameliorate or compensate cellular and circuit deficits in brain disorders.

## AUTHOR CONTRIBUTIONS

Z.J.H. and A.P. conceived the study. Z.J.H. organized and supervised the study. A.P. performed scRNAseq experiments. A.P. and R.R. performed mRNA in situ and immunohistochemistry experiments. A.P., Z.J.H., J.G., M.C. analyzed the data. M.C. and J.G. developed MetaNeighbour and performed DE, AUROC analysis. Z.J.H. and A.P. wrote the manuscript with contributions from J.G. and M.C.

## ACKNOWLEDGMENT

We acknowledge Dr. Michael Wigler for sharing Unique Molecular Identifier strategy and barcodes (varietal tags), Jude Kendall for programming support, Drs. David Spector and Nour El-Amine for help with super-resolution microscopy and Priscila Wu for technical assistance.

We thank Drs. Liqun Luo, Chris McBain, Tony Zador and Linda Van Aelst for valuable comments and suggestions on the manuscript. This work was supported by NIH 5R01MH094705-04, R01MH109665-01 and CSHL Robertson Neuroscience Fund to Z.J.H. A.P. was supported by a NARSAD Postdoctoral Fellowship. J.G. and M.C. were supported by a gift from T. and V. Stanley.

## Supplementary figures

**Figure S1.**
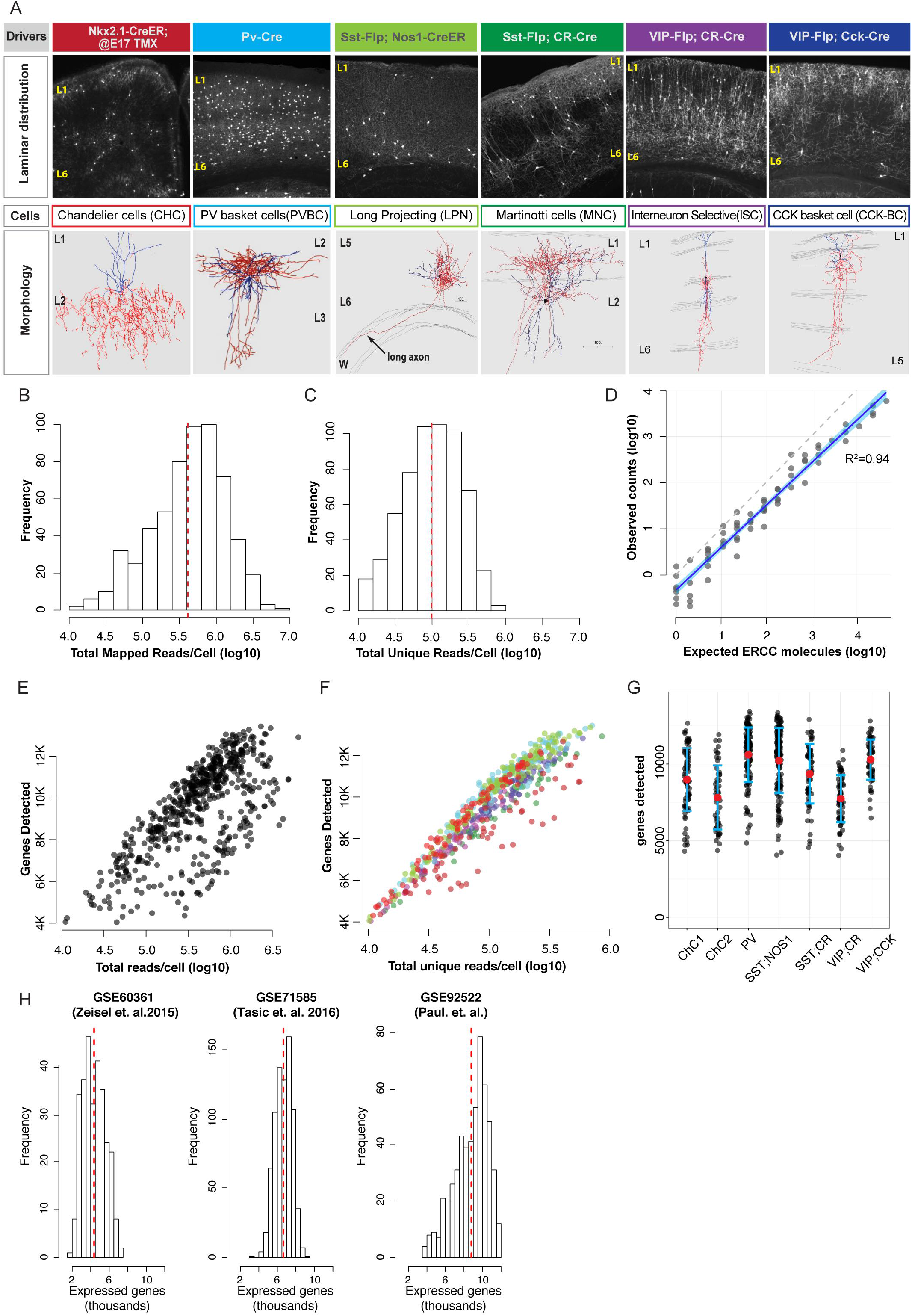
Transcriptomic analysis of cortical GABAergic ground truth populations. **(A)** Combinatorial genetic labeling of cortical GABAergic GTPs. Intersectional, lineage and birth time dependent labeling (top row) of 6 GTPs and bona fide types with characteristic laminar distribution (middle row). Representative single cell reconstruction depicted characteristic morphology of bona fide cell type in each GTP (bottom row). **(B)** Histogram of total mapped reads per single cell with a median read depth of 4.0x10^5^ reads per cell. **(C)** Histogram of unique reads per single cell with a median of 1.0x10^5^ counts. **(D)** Plot of ERCC observed unique reads versus the numbers of molecules expected for each species from the ERCC cocktail shows linearity and slope close to 1; unity line shown as dotted grey line, blue shaded region shows 95% confidence interval. **(E)** Number of genes detected in single cells versus total mapped reads and **(F)** unique reads shows that gene detectability is correlated to read counts but there are no gene detection bias towards any of the GTPs (color code as in A). **(G)** Genes detected across GTPs range between 7.5 – 12 thousands (median). **(H)** Higher levels of genes detected in single cells compared to GSE60361 (Zeisel et al., 2015) and GE71585 (Tasic et al., 2016) dataset.

**Figure S2.**
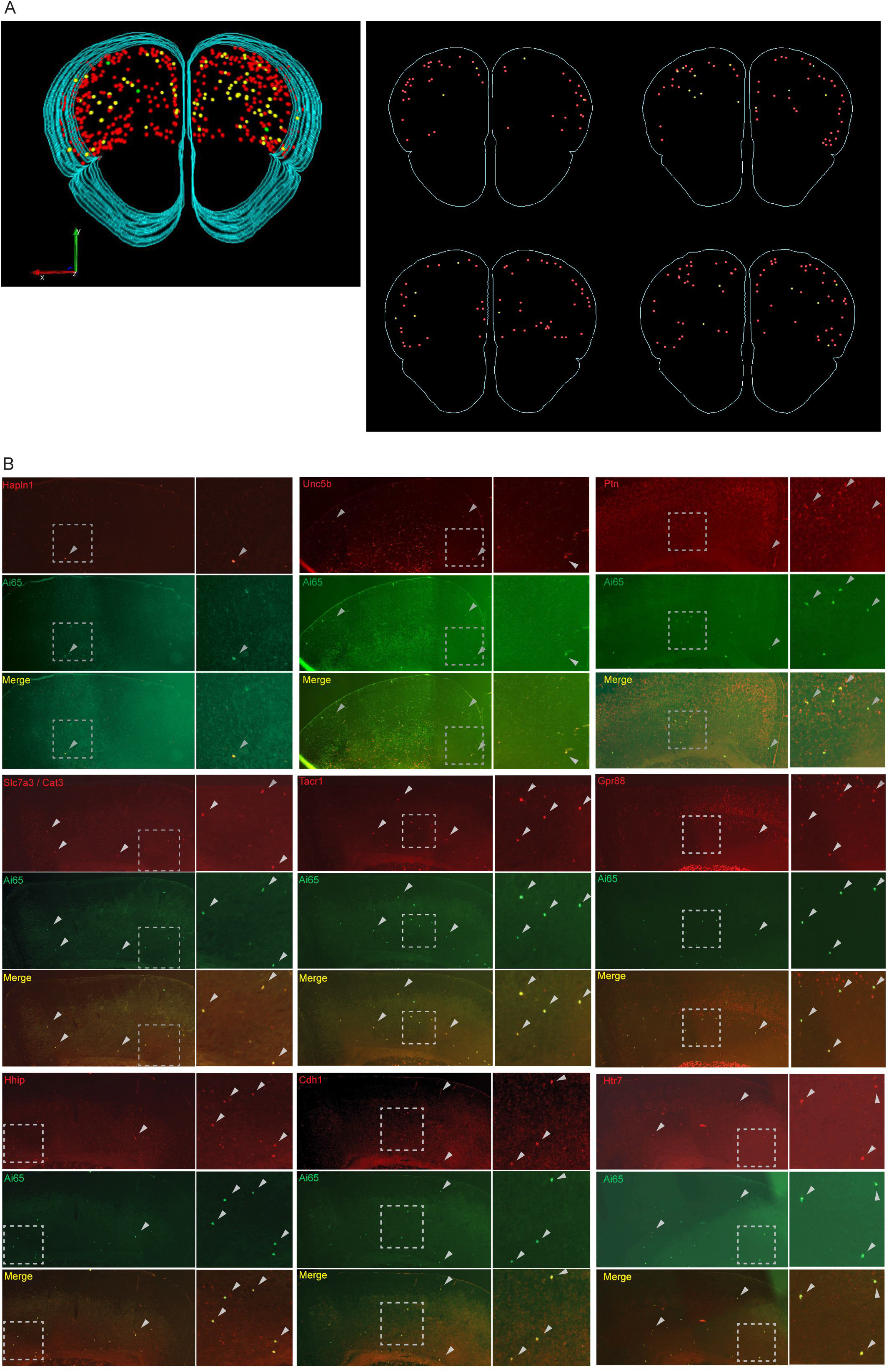
Fluorescent double in-situ in brain tissue confirms co-localization of select enriched transcripts in GTPs. **(A)** Left: Partial 3D reconstruction render from serial coronal sections (spanning 288um, rostro-caudal) of mouse forebrain showing double mRNA in situ of CHC enriched transcript Pthlh (red), Ai14/tdTomato (green) and co-localization (yellow). Only a subset of CHCs were labeled in *Nkx2.1-CreER*;Ai14 mice by tamoxifen induction at E17. Right: representative sections from 3D render. 136/143 tdTomato^+^ (~95%) co-express Pthlh. Laminar distribution of Pthlh signal was similar to those reported for ChC distribution pattern (Taniguchi et. al. 2013). **(B)** Representative single images from double RNA in situ of Ai14/tdTomato (green) shows colocalization (yellow) with select transcripts enriched in CHC2 and (Unc5b, Hapln1) SST;NOS1 cells (Ptn, Slc7a3, Tacr1, Gpr88, Hhip, Cdh1 and Hrt7). Dotted box represents area in higher magnification, arrowheads indicate co-localization.

**Figure S3:**
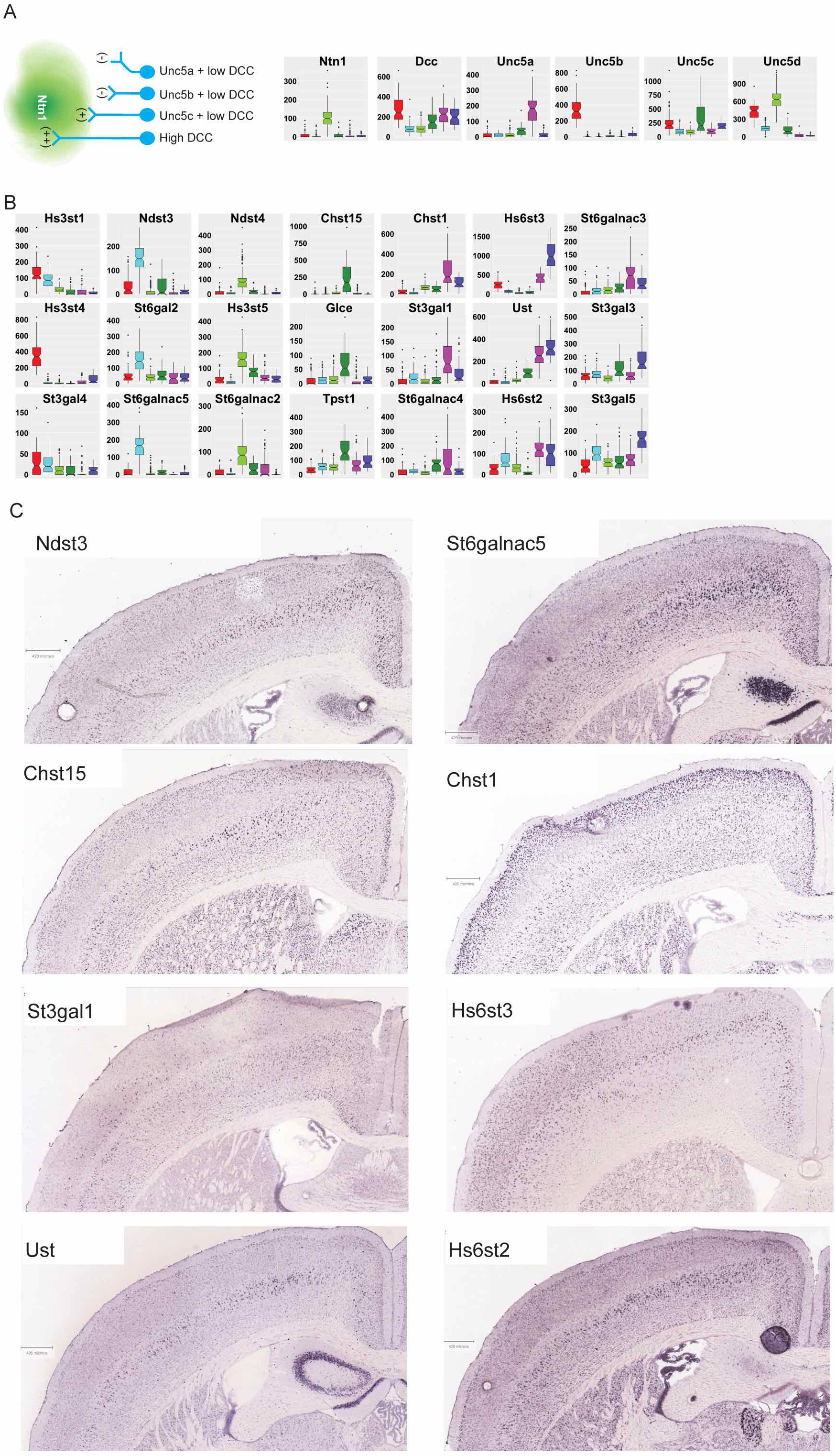
Differential expression of netrin-unc5 and carbohydrate modifying enzymes among GTPs. **(A)** Schematic depiction of known attractive (+) and repulsive (-) interactions between the ligand Netrin (Ntn1) and its receptors - Unc5 family members and DCC (Kolodkin et al., 2011). Boxplots show highly specific expression of Ntn1 and Unc5 family members among GTPs. **(B)** Boxplots of CAM modifying enzymes Sulfo- and Sialyl-transferases enriched in individual GTPs. **(C)** Representative in situ images from Allen Brain Atlas showing distinct laminar distribution patterns of some carbohydrate modifying enzymes in coronal brain sections. Many of these patterns are characteristic to the distribution of subsets of GABAergic neurons.

**Figure S4:**
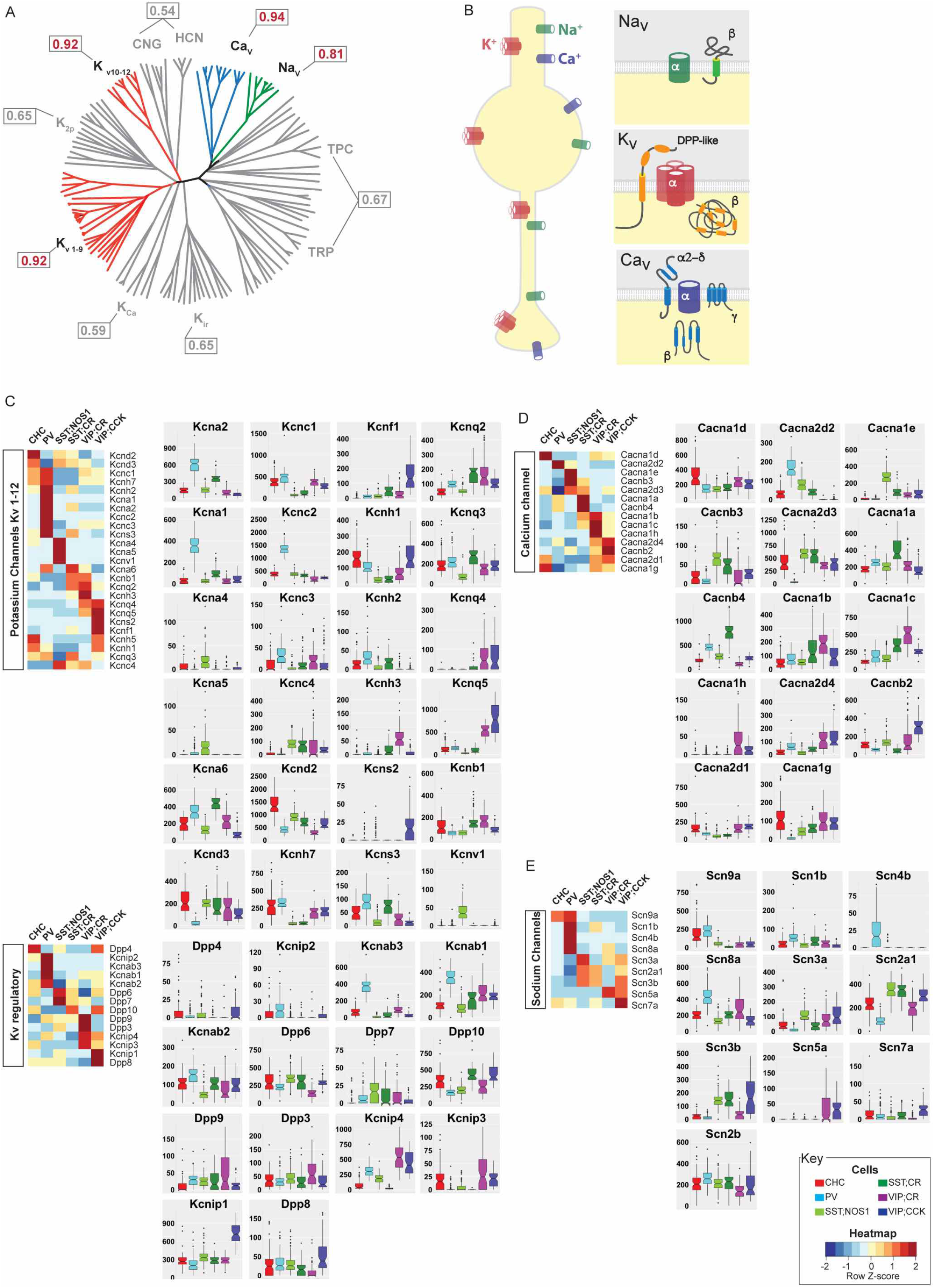
Differential expression of voltage-gated ion channels among GTPs. **(A)** An unrooted cladistic representation drawn using minimum evolution analysis of 143 members of structurally related voltage-gated ion channels based on their amino acid sequence of the minimal pore regions adopted without structural manipulations (modified from Yu and Catterall, 2004). Only three of these families, Kv, NaV and CaV are highly discriminative of GTPs as shown by their AUROC scores (boxed, red text shows high scoring families). **(B)** Schematic of the subcellular distribution of voltage gated ion channels and their subunit compositions of pore forming and accessory regulatory proteins. **(C)** Heatmap and selected boxplots showing differential expression of Kv channels and their regulatory subunits among GTPs. Fast-spiking PV basket cells express the highest levels and diversity of Kv channels. **(D)** Heatmap and selected boxplots showing differential expression of CaV channels. **(E)** Heatmap and selected boxplots showing differential expression of NaV channels. Fast-spiking PV basket cells express the highest levels and diversity of Nav channels

**Figure S5:**
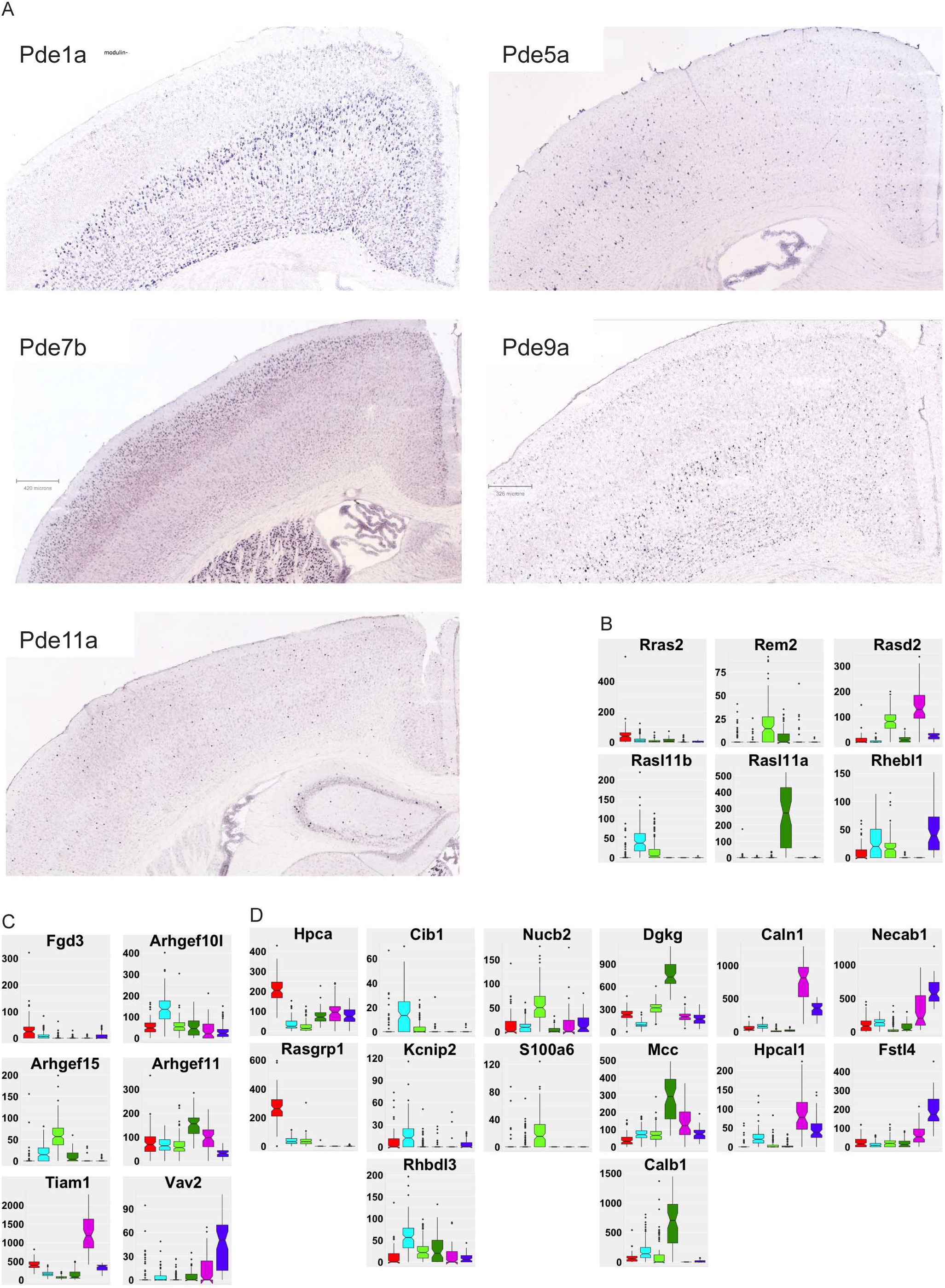
**(A)** Selected RNA in-situ images from Allen Brain showing that multiple phosphodiesterases are expressed in restricted neuronal populations in adult mouse cortex with patterns characteristic to GABAergic interneurons. **(B-D)** Boxplots for Ras (B), Rho-GEF (C), and EF-hand/CaBP (D) family members among GTPs.

**Figure S6:**
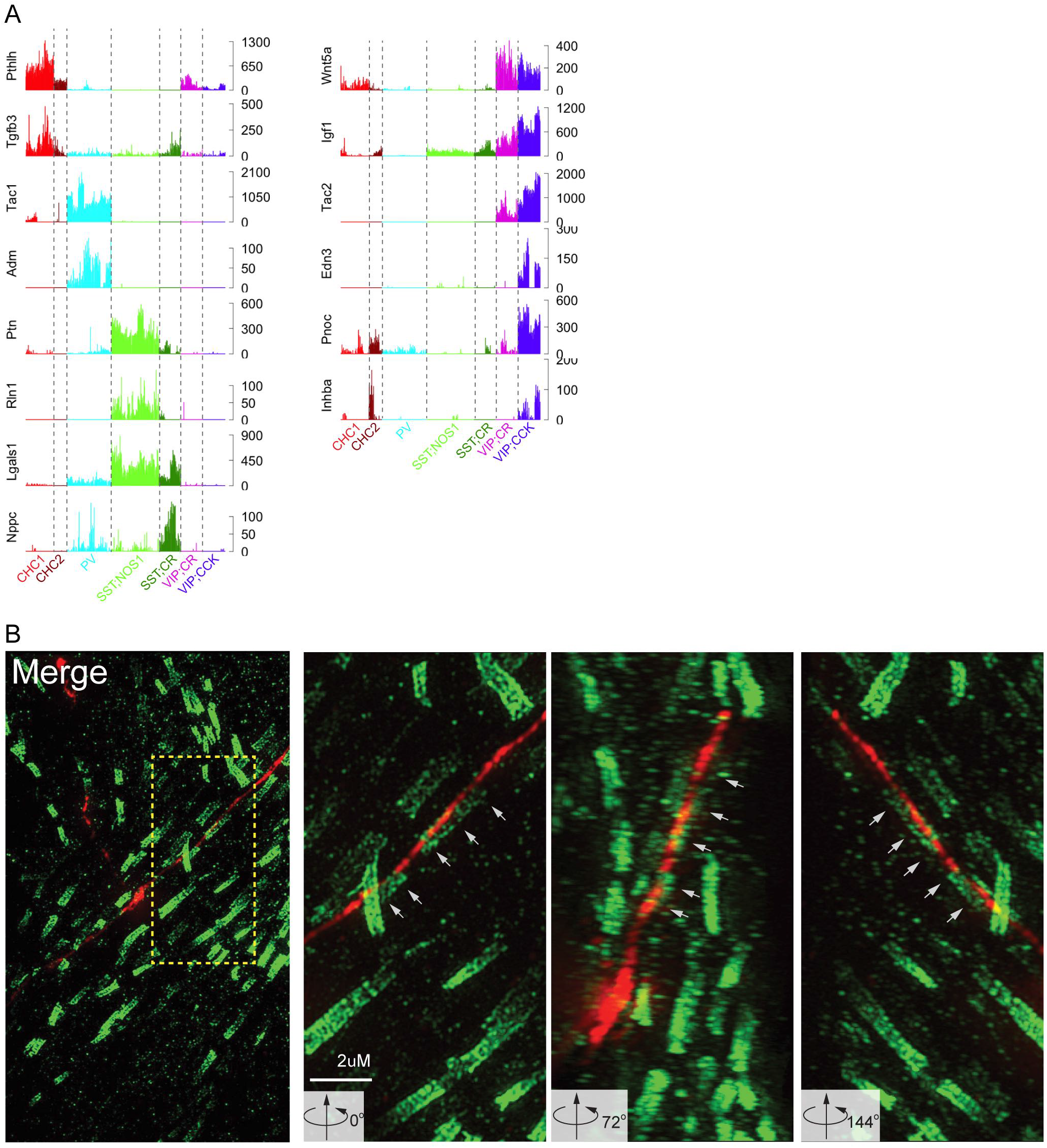
SST;NOS1 axons are myelinated. **(A)** Barplot of neuropeptides and endogenous ligands expressed in ON/OFF pattern in individual cells of each GTP (colors). **(B)** Left: 63X image shows SST;NOS1 axons traversing deep cortical layers, among a field of myelin nodes labeled by CASPR immunostaining. Dotted box is enlarged in the right 3 panels, which show 3D rendering of higher magnification images and coaxially apposed CASPR signal over tdTomato-labeled axons of SST;NOS1 at three angular rotations (white arrows).

**Figure S7:**
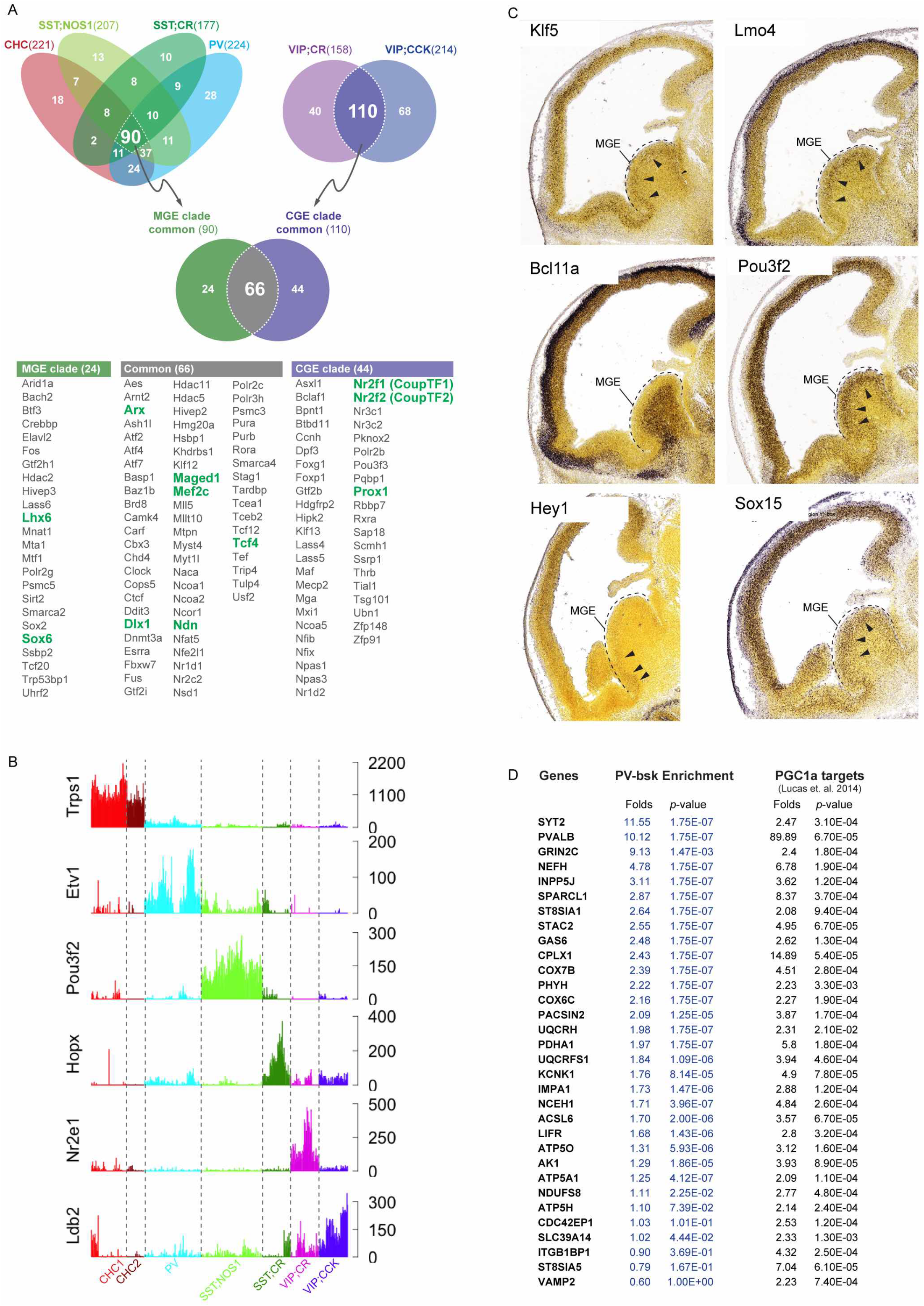
Transcription factor profiles among GTPs and PGC1α transcription network. **(A)** Top: Venn diagram of all TFs expressed in each GTP; TFs were assigned to a set if it is expressed in ≥75% of cells within a GTP at a level of ≥30uTPM in single cells. Common TFs in MGE and CGE derived GTPs are further intersected to generate a list of common GABAergic TFs. Bottom: Table of TFs in each set, green fonts indicate TFs known to be expressed as computed from the Venn diagram. **(B)** Barplots of TFs expressed in ON/OFF pattern in individual cells of GTPs **(C)** mRNA in situ Images from Allen Developmental Mouse Brain Atlas at E13.5 show expression of transcripts in the MGE that are expressed in PV<SST and SST;CR and SST;NOS1 from Fig-7F and G. **(D)** Table showing that >75% of PGC1α targets identified by Lucas et. al. 2014 are also significantly (p<1x10^-05^) enriched in PVC transcriptome data.

## List of Supplementary Tables

Table S1. DE table of genes in Fig1D, FC>4, uTPM>50, q value<5.0 X 10^-4^, batches>80%

Table S2. DE table of high performance CAMs in Fig3E

Table S3. AUROCs values for ~3888 GO gene sets

Table S4. AUROC scores of ~450 HGNC gene families from 3 studies (Paul et. al., Tasic et. al. 2016 and Zeisel. et. al. 2015)

Table S5. AUROC scores of custom gene sets used

## Methods

Contact for reagent and resource sharing

Dr. Z. Josh Huang

### Animals

Nkx2.1-CreER, Pv-ires-Cre animals were bred separately to Ai14 reporter to label CHC and PVBC in the cortex. CHCs were enriched in frontal cortex with tamoxifen induction at E17.5. Intersectional labeling of GTPs were achieved by breeding (a) Sst-Flp, Nos1-CreER, (b) Sst-Flp, CR-Cre, (c) VIP-Flp, CR-Cre and (d) VIP-Flp, CCK-Cre separately to Ai65 intersectional reporter that will label cells with tdTomato only when both the lox-Stop-lox and Frt-STOP-Frt cassettes are excised (see supplementary Fig-S1). Mice were bred and maintained according to animal husbandry protocols at Cold Spring Harbor Laboratory (Institutional Animal Care and Use Committee reference number 16-13-09-8) with access to food and water ad libitum and 12 h light-dark cycle.

### Manual cell sorting

To isolate individual RFP-labeled GABAergic neurons, we microdissected motor and somatosensory cortical slices from fresh brain tissues of mature (6 weeks old) mice, generated single cell suspension and manually purified single RFP-labeled cells (Paul et al., 2012; Sugino et al., 2006). Brains were sectioned at 300 μm thickness using a cooled stage vibratome (Microm, Model HM360) with circulating oxygenated artificial cerebrospinal fluid. Sections were blocked in AP5, CNQX, and TTX cocktail to prevent excitotoxic cell death and then treated with mild protease (Fraction IV protease Streptomyces, Sigma Cat#P5147-5G). Brain regions of interest were microdissected and triturated to dissociate the cells. Dissociated cells were put into in a Petri dish in low density for optimal cell-cell separation and single RFP-positive cells was collected using patch pipette capillary and dispensed individually into separate single tubes prefilled with RNAseOUT (Invitrogen), ERCC spike-in RNAs in 1:400 K dilution, sample specific RT primers for a total of 1 μL volume. Process was repeated to collect 32-64 cells in one manual cell sorting session. Cells were flash frozen in liquid nitrogen and stored at – 80 °C until processed. Patch pipette was single use only and fresh pipettes were used for every single cell collected.

### Linear RNA amplification, Illumina library prep, and sequencing

Single cell mRNAs were converted to cDNAs through polyA primers (Eberwine et. al. 1992) containing a sample barcode and unique molecular identifiers (UMIs). We employed two rounds of in vitro transcription amplification (Hashimshony et al., 2012; Jaitin et al., 2014) followed by Illumina TrueSeq protocol to construct RNAseq libraries.

Custom T7-polyA primers with were designed containing additional 9bp error correcting sequences for identifying single cells (sample barcode) and 10bp random nucleotide sequences (UMI/varietal tag) to label each mRNA molecule amplified with a unique barcode. The UMI allows for elimination of reads containing duplicate tags for the same mapped sequence and only tally up the total unique tags of all mapped sequence to a coding sequence. This primer also contained a 26bp flanking RA5 adapter sequence needed for downstream illumina cDNA library step, which eliminates a rate limiting enzymatic 5’ ligation step of cDNA preparation increasing efficiency (Hashimshony et. al. 2012).

RNA was linearly amplified by T7 RNA polymerase using two rounds of in-vitro transcription (MessageAmp-II kit Life Technologies) according to the manufacturer’s recommended protocol with some modifications. Cells were lysed by repeated heating to 70C and snap cooling to 4C and first strand synthesis was carried out at 42C for 2hrs with first strand buffer, dNTP mix, RNase inhibitor and ArrayScript enzyme. Second strand synthesis was done at 16C for 2hrs with second strand buffer, dNTP mix, T4 DNA polymerase and RNAseH. cDNA was purified using columns and first round IVT was performed at 37C for 14hrs to make aRNA. For the second round of linear amplification column purified aRNA from first IVT underwent another first strand synthesis at 42C for 2hrs, followed by RNaseH digestion at 37C for 30mins and another second strand synthesis at 16C. The resulting double stranded cDNA underwent a final second IVT step at 37C for 14hrs to make aRNA. These linearly amplified aRNA products now carried the 3’-end of the polyA transcripts for mapping to coding regions plus the sample barcode to indicate which GTP it came from, UMI sequence for counting unique cell-endogenous parent mRNA molecules and one of the flanking sequence (RA5 adapter) for Illumina sequencing. Second round aRNAs were fragmented chemically using NEBNext^®^ Magnesium RNA Fragmentation Module (Cat#E6150S), column purified using RNA MinElute (Qiagen) for final Illumina cDNA library preparation steps.

cDNA library was generated using Illumina TruSeq small RNA kit (Cat#RS-200-0012) and only 3’-adapter (RA3) need to be ligated enzymatically using truncated T4 RNA ligase (NEB M0242) on to the fragmented aRNA and the 5’ ligation step for RA5-adapter was skipped (Hashimshony et. al. 2012). Adapter ligated fragmented aRNA was reverse transcribed using SuperScriptIII reverse transcriptase (Invitrogen, USA) and PCR enriched using TruSeq indices (for multiplexing) for no more than 7–11 cycles. The resulting library was size-selected using SPRISelect magnetic beads (Agencourt) to select 350-450bp fragments and paired-end sequenced for 101bp in Illumina HiSeq. No more than 32 single cells were run in one lane of HiSeq2000 generating on average ~180-200 million reads per lane.

### Mapping and tag counting

As in our previous work (Crow et al, 2016), Bowtie (v 0.12.7) was used for sequence alignment of read2 (polyA primed) to the mouse reference genome (mm9), while read1 sequences were used for UMI (varietal-tag) counting. A custom python script was used Bowtie (v 0.12.7) was used for sequence alignment of read2 (polyA primed) to the mouse reference genome (mm9), while read1 sequences were used for UMI (varietal-tag) counting. Multiple reads to the same gene with the same tag sequences were rejected and only counted as one, such that only mapped sequences with unique tags were retained and tallied for each mRNA for each cell.

We obtained ~ 4.8X10^5^ (median, Avg=6.9x10^5^) mapped reads per cell, each containing ~1.0x10^5^ (median, Avg=1.4x10^5^) unique reads that typically detected on average ~10,000 genes (range ~7,500 to ~12,000 genes median), with >95% of the single cells detecting >6,000 genes (Figure S1B-C). In each single cell ERCC spike-in RNA (Life Technologies) were used as internal controls, for which the absolute number of molecules that are added to sample can be calculated; this gave a linear relationship of input-output measures with a slope of 0.92 and adjusted R^2^=0.96 (Figure S1D). Following quality control screen, we obtained high depth transcriptome of ~584 cells from the 6 GTPs (Figure S1D). This unique dataset thus contains high-resolution transcriptomes of phenotype-defined cortical GABAergic GTPs.

For any given gene the absolute unique counts were normalized to the total unique counts across all genes in a single cell and are expressed as unique Transcripts Per Million (uTPM). To determine differential gene expression (DE) and calculate fold-change, gene-wise Fisher’s meta-analytic p-value was calculated on these normalized gene expression values without further batch effect correction.

### Fisher's meta-analytic DE

To assess differential gene expression across GTPs, we took advantage of the replicate batches within each type and performed a meta-analysis across replicates based on non-parametric statistics. Briefly, for each GTP we performed one-tailed Mann-Whitney tests between individual batches within a cell type against all cells outside of that cell type. To ensure that significance would arise from replication rather than extreme p-values, prior to meta-analysis with Fisher’s method, p-values at FDR<=0.05 for an individual test were set to the maximum p-value meeting that criterion. Finally, meta-analytic p-values were FDR corrected. Differentially expressed gene sets were defined by FDR adjusted p-value <0.05 having log2 fold change >2

### MetaNeighbor (AUROC calculation)

To measure GTP identity we use the MetaNeighbor method as described in our companion paper (Crow et. al. 2017). In brief, MetaNeighbor requires the input of a set of genes, an expression matrix and two sets of labels: one set for labeling each experiment, and one set for labeling the cell types of interest. Here, each batch was treated as an “experiment”, and we aimed to measure the replicability of cell identity across batches. Cell-type labels are held back from one experiment at a time and then predicted based on the others, to determine which gene sets functionally characterize cells across technical variation. For each gene set being used to evaluate a given cell-type, the method generates a network based on the Spearman correlation between all cells across the genes within the set. The correlation is rank standardized to provide network weightings between each pair of cells, and then a neighbor voting predictor scores cells as possessing a given annotation. The score is calculated as the sum of a given cell’s connectivity weighting to neighbors possessing a given cell annotation. For cross-validation, we permute through all possible combinations of leave-one-batch-out cross-validation, and report the degree to which cells of the same type are recovered as the mean area under the receiver operator characteristic curve (AUROC) across all folds. To improve speed, AUROCs are calculated analytically:

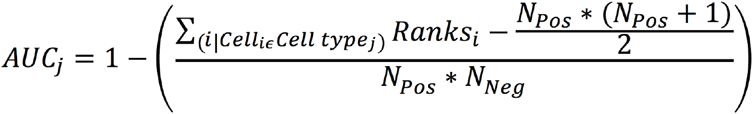

where “Ranks” are the ranks of the hidden positives, N_pos_ is the number of true positives, and N_Neg_ is the number of true negatives. A Github repository containing R scripts and parsed data can be found online https://github.com/maggiecrow/MetaNeighbor.

Single cell mRNAs were converted to cDNAs through polyA primers containing a sample barcode and unique molecular identifiers (UMIs). We employed two rounds of in vitro transcription amplification (Hashimshony et al., 2012; Jaitin et al., 2014) followed by Illumina TrueSeq protocol to construct RNAseq libraries. Through next generation sequencing, we obtained ~ 4.8X10^5^ (median, Avg=6.9x10^5^) mapped reads per cell, each containing ~1.0x10^5^ (median, Avg=1.4x10^5^) unique reads that typically detected on average ~10,000 genes (range ~7,500 to ~12,000 genes median), with >95% of the single cells detecting >6,000 genes (Figure S1B-C). In each single cell ERCC spike-in RNA (Life Technologies) were used as internal controls, for which the absolute number of molecules that are added to sample can be calculated; this gave a linear relationship of input-output measures with a slope of 0.92 and adjusted R^2^=0.96 (Figure S1D). Following quality control screen, we obtained high depth transcriptome of ~584 cells from the 6 GTPs (Figure S1D).

This unique dataset thus contains high-resolution transcriptomes of phenotype-defined cortical GABAergic GTPs.

### RNA double in-situ and imaging

RNA double in-situ was performed using Quantigene ViewRNA tissue ISH (Affymetrix, USA) following manufacturer's recommended protocol. Fresh unfixed brain tissues were frozen in OCT blocks using dry-ice isopentane slurry. Brains can be stored in -80C until cryosectioning. Cryosectioning was done on Leica cryotome at 12um thickness, and sections collected on charged glass slides. Custom and off-the shelf branched-DNA oligo ISH probes were designed and synthesized by Affymetrix Quantigene ViewRNA. Sections on slides were postfixed just prior to ISH, and in-situ steps were followed according to manufacturer’s recommended protocol. For dual signal detection QuantiGene Type-1 and Type-6 probes were used. Fluorescent signals from Type-1 and Type-6 ISH probes were imaged on tile-scanning mode using Perkin Elmer spinning disk confocal at 10X magnification and auto-stitched using Volocity software. Stitched images were exported as TIFFs for further processing and adjustments to brightness and contrast in FIJI (Fiji is just ImageJ) and assembled in Adobe Illustrator.

### Super-resolution microscopy

Super resolution images were acquired with GE Healthcare OMX V3 system using: 488 and 593 nm solid state lasers; UPlanS Apochromat 100×1.4 NA objective lens (Olympus); 2 EM-CCD cameras (Cascade II 512, Photometrics). 3D structured illumination images were reconstructed with SoftWoRx^®^ 6.5.2 software. 3D rendering was performed using Imaris (Bitplane) 7.6.5. exported as TIFF, processed in FIJI and assembled in Adobe Illustrator.

### Ethics

Mice were bred and maintained according to animal husbandry protocols at Cold Spring Harbor Laboratory (Institutional Animal Care and Use Committee reference number 16-13-09-8).

### Glossary of terms

- Neighbor voting – A method to classify cells into known types based on the similarity of the gene expression profiles for a given gene set.
- Cross-validation – A method to estimate how well the results of an analysis will generalize. The main purpose of cross-validation is to avoid overfitting to a particular dataset, or in this case, library batch. There are many ways to implement cross-validation; in this study we use a batch-stratified cross-validation. In this analysis, we hide labels from one batch at a time, making predictions as each label is hidden, until we have tested all of the batches within a particular cell type.
- Performance – The metric used to quantify cell identity. This refers to the area under the receiver operating characteristic curve (AUROC).

## SUPPLEMENTAL TEXT

### 1. Differential expression of cell adhesion molecules and carbohydrate modifying enzymes among GTPs suggests large capacity for cell surface and extracellular matrix labels

#### 1a. Cell adhesion molecules

Among IgCAMs, Kitl, Kirrel, JAM2, 3, ICAM5 are each highly enriched in a specific subpopulation (Figure 3F). Other differentially expressed cell adhesion systems include: non-clustered protocadherins, major cadherins, receptor protein tyrosine phosphatases, semaphorin-plexin group (Figure 3F).

#### 1b. Carbohydrate modifying enzymes that diversify proteoglycans and glycoproteins

In addition to adhesion proteins serving as membrane labels, cell surface is extensively decorated by a large variety of carbohydrates presented by glycoproteins (many are glycosylated CAMs) and especially by proteoglycans along the membrane extending into the extracellular matrix. Among major groups of proteoglycans, heparan sulfate proteoglycans cooperate with CAMs to mediate cell-cell and cell-matrix interactions (Lee and Chien, 2004; Smith et al., 2015). On the other hand, chondroitin sulfate proteoglycans inhibit neurite growth and plasticity and form lattice like perineuronal nets (PNN) around subsets of neurons, including cortical PV basket cells (Miyata and Kitagawa, 2016a). The progressive postnatal formation of PNNs promotes the functional maturation of PV cells and regulates the critical period of neural plasticity in visual cortex (Pizzorusso et al., 2002). Interestingly, different neuron types may be surrounded by chondroitin sulfates of different structures that accumulate different proteins type (Matthews et al., 2002). It is unknown whether different GABAergic neurons generate unique blend of extracellular matrix such as distinct carbohydrate coats (i.e. a sugar code).

Proteoglycans consist of long polysaccharide chains covalently linked to one of a small number of core proteins (Miyata and Kitagawa, 2016b; Sarrazin et al., 2011). The extremely large molecular diversity of proteoglycans derives from the modification patterns, especially sulfation patterns, of sugar residues along the carbohydrate chain and at specific positions. These modifications are carried out by a large family of sulfotransferases in the Golgi apparatus, each catalyzes sulfation at specific carbon position of the sugar residue ring. Thus it is possible that a cell might express a particular combination of sulfotransferases to produce specific sulfation patterns for its proteoglycan repertoire (Lee and Chien, 2004). Indeed, developmental changes in the expression of two different sulfotransferases in PV interneurons alter the ratio of 4-sulfation/6-sulfation ratio of CSPG and regulate the maturation of PNN and PV function (Miyata et al., 2012). In addition to sulfation, glycoproteins can also be coated by sialic acid onto their carbohydrate chains, catalyzed by a family of sialyltransferases (Schnaar et al., 2014). However, the cellular expression of sulfotransferase repertoire is largely unknown. We discovered that both the sulfotransferase and sialyltransferase families are differentially expressed among GTPs, with high AUROC scores (0.88 and 0.85, respectively). Strikingly, six sulfotransferase are highly specific to different GTPs (Figure S3B): Hs3st4 to ChC, Chst15 to SST/CR, Chst1 to VIP/CR, Hs3st5 to SST/NOS; Hs3st1 is enriched in ChC and PV cells and Hs6st3 is enriched in VIP/CCK cells. Differential combinations of these carbohydrate modifying enzymes may generate a characteristic repertoire of sugar-decorated proteoglycans that customize the cell coat and extracellular matrix.

## 2. Differential expression of transmitter and modulator receptors shapes input propertie: of GTPs

### Ionotropic GABA receptors (GABAARs)

Martinotti cells are considered “master regulators” that innervate most other cell types, including the distal dendrites of pyramidal neurons (Jiang et al., 2015). Yet they avoid themselves and receive relatively few inhibitory inputs, with the prominent exception of inhibition from VIP cells (Jiang et al., 2015; Pfeffer et al., 2013). With highly limited subunit repertoire, we infer that SST/CR cells most likely receive VIP cell inputs through α3β1/3γ3 type GABA_A_Rs.

Another major inhibitory mechanism is the dis-inhibitory module represented by interneuron selective VIP cells, which target predominantly Martinotti and to a less extent PV cells, but not themselves (Jiang et al., 2015; Pfeffer et al., 2013; Pi et al., 2013). They receive relatively few local excitatory and inhibitory inputs, but are innervated by PV and SST cells (Jiang et al., 2015; Staiger et al., 1997). It is possible that α1-containing GABAARs mediate PV cell input whereas α3-containing GABA_A_Rs mediate Martinotti cell input.

## 3. Differential expression of signaling proteins in calcium, cyclic nucleotide and small GTPase 2^nd^ messenger pathways customizes intracellular signaling in GTPs

### 3a. Ca2+ binding proteins likely shapes spatiotemporal dynamics of Ca2+ signaling

Many ligand-gated ion channels conduct Ca2+, a ubiquitous and versatile 2nd messenger, to trigger intracellular signaling (Brini et al., 2014; Burgoyne and Haynes, 2015). The spatial changes of intracellular Ca2+ are often confined to micro- and nano-domains (Eggermann et al., 2011), as excess “free” Ca2+ ions is cytotoxic; and the temporal dynamics of Ca signal ranges from micro-seconds to minutes, and from pulsatile to oscillatory (Bading, 2013; Dupont, 2014). These exquisite spatiotemporal patterns confer the specificity and potency of Ca2+ signaling through a large variety of Ca2+-regulated enzymes and effectors (Brini et al., 2014). To a great extent, spatiotemporal Ca2+ dynamics are shaped by a large set of Ca2+-binding and signaling proteins (CaBPs) (Burgoyne and Haynes, 2015). The mouse genome encodes ~170 EF-hand containing CaBPs with distinct binding affinities, kinetics and subcellular localization. Beyond the well characterized CaBPs as GABAergic markers (e.g. PV, calretinin, calbindin) (Kubota et al., 2011), the expression of most CaBPs in different neuronal cell types are unknown. We found that each GTP expresses a set of ~5-8 different CaBPs (Figure 5D). Many of these CaBPs are in fact signaling proteins (e.g. Rasgrp1 in ChCs). These results suggest that differential expression of multiple Ca2+ binding and signaling proteins might shape distinct spatiotemporal dynamics and the specificity of Ca2+signaling among GTPs.

### 3b. Adenylyl cyclase and phosphodiesterase isoforms may shape distinct cAMP signaling properties

GPCRs signal through G proteins, many of which engage cAMP - the archetypical 2nd messenger pathway. cAMP activates protein kinase A (PKA) which regulates effector proteins through phosphorylation. The synthesis, degradation and spatiotemporal dynamics of cAMP are stringently regulated at each step (Halls and Cooper, 2011). First, different Gα subunits either activate or inhibit different isoforms of adenylyl cyclases (ACs), which catalyze rapid cAMP synthesis; the activities of Gα subunits are tightly controlled by a family of regulators of G protein signaling (RGS) (Gerber et al., 2016). In parallel, the equally rapid degradation of cAMP is mediated by a large family of phosphodiesterases (PDEs) with distinct catalytic and regulatory properties (Maurice et al., 2014). As brain tissues contain 10fold greater PDE than AC activity (Schmidt, 2010), the intricate coordination of specific AC and PDE activities tightly controls the spatiotemporal changes of cAMP concentration (McCormick and Baillie, 2014). Importantly, different isoforms of ACs and PDEs localize to various subcellular compartments and further assemble into specific signaling complexes through binding to designated scaffolding proteins (e.g. A kinase adaptor proteins, or AKAPs), which recruit appropriated PKA isoforms and their particular target effectors (Edwards et al., 2012). These signaling complexes thus achieve exquisite specificity by presenting particular “flavors” of cAMP signal to specific effectors through physical proximity (McCormick and Baillie, 2014). The mouse genome contains 13 Gα, 7 Gβ, 12 Gγ, 23 RGSs, 9 ACs, 22 PDEs, 25 AKAPs, and 13 PKA subunits. The extent to which members of these signaling protein families are customized for specific neuronal cell types are unknown. Our computation screen and analysis revealed highly coordinated differential expression across these families of signaling proteins among GTPs.

### 3c. cGMP signaling modules in SST/nNOS and ChC

In contrast to cAMP, which serves as a ubiquitous 2nd messenger for vast number of extracellular ligands through hundreds of GPCRs, cGMP signaling in the brain is predominantly if not specifically triggered by nitric oxide (NO) (Lucas et al., 2010). NO is synthesized by neuronal nitric oxide synthase (nNOS) from L-arginine, which is acquired by neurons through specific amino acid transporters (Slc7a1-3) (Friebe and Koesling, 2003). In mature cortex, nNOS is expressed in subsets of GABAergic neurons, with high levels in a small set of SST+ long projection cells (LPCs, also type I nNOS cells) and much lower levels in several other populations (type II nNOS cells) (Perrenoud et al., 2012; Taniguchi et al., 2011). The major target of NO is the soluble form of guanylyl cyclase which catalyzes cGMP production. cGMP modulates the activity of cyclic nucleotide gated channels and PDE2/3, and engages protein kinase G (PKG) to regulate downstream effectors through phosphorylation (Friebe and Koesling, 2003). Although this general scheme is well established in brain tissues, whether and how NO and cGMP signaling is differentially implemented in different neuronal cell types is far from clear. We have found highly distinct mode of cGMP signaling among GTPs and discovered signaling modules in two bona-fide cell types (Figure 5A, E).

As the major effectors of PKG are ion channels, we screened through ion channels that are enriched in LPCs and ChCs (Figure S4) and searched for potential PKG targets by literature curation of their regulation by phosphorylation. We found at least two members of the Trp (transient receptor potential) channels and BK-type potassium channels that are differentially enriched in these two cell types and have been shown to be NO and PKG targets. The large conductance Ca- and voltage-activated potassium channels (BK-type) consist of a α1 core subunit and β auxiliary subunits. Channel activity is stimulated by PKG phosphorylation of the pore-forming α1 subunit (KCNMA1) (Alioua et al., 1998; Kyle et al., 2013; Zhou et al., 2001). We found co-expression of α1 and β2 in ChCs, which assemble the fast activated and inactivating form (Wang et al., 2014), consistent with their fast electrophysiological properties. In contrast, α1 and β4 are enriched in LPCs, which assemble the slow activated and non-inactivating form (Wang et al., 2014), consistent with their multiple slow forms electrophysiological properties. In the Trp family, Trpc6 is 6 times more permeable to Ca2+ than to Na+ and can activate transcriptional pathways involving calmodulin kinase IV (CAMKIV) and cAMP response element-binding protein (CREB) (Dietrich and Gudermann, 2014), while Trpc5 exhibits slightly higher permeability to Ca2+ over Na+ (Zholos, 2014). While Trpc6 is a PKG target (Takahashi et al., 2008) and is enriched in LPCs (Figure 5F), Trpc5 is directly activated by NO-mediated cysteines S-nitrosylation (Yoshida et al., 2006) and is enriched in ChC.

### 3d. Differential expression of Ras and Rho small GTPases

In addition to transmitters, modulators and hormones, cortical neurons respond to a diverse set of membrane bound or diffusible protein ligands that mediate cell-cell contacts and signaling through receptor tyrosine kinases (RTKs; Figure 5A). A key step of RTK signaling is mediated by a large set of Ras superfamily small GTPases, which play similar roles as classic 2nd messengers to activate multiple, specific kinase cascades that engage effectors (Alberts, 2014; Colicelli, 2004). As highly versatile molecular switches, small GTPases are activated by guanine nucleotide exchange factors (GEFs) and inactivated by GTPase activating proteins (GAPs) (Cherfils and Zeghouf, 2013). Within the Ras superfamily, only the Ras and Rho families relay membrane receptor signals, each family is regulated by a designated set of GEFs and GAPs (Cook et al., 2014; Vigil et al., 2010). Upon ligand activation, tyrosine phosphorylation in RTK cytoplasmic domain recruits specific adaptor proteins, which further assemble GEFs and GAPs to regulate Ras and Rho GTPases and engage kinase cascades (Alberts, 2014). Prominent among diverse effectors of the RTK pathways are transcription factors, which regulate gene expression (Ye and Carew, 2010), and cytoskeleton proteins that regulate cell shape, motility, adhesion and intracellular transport (Soderling, 2014).

The mammalian genome contains ~60 RTKs, over 90 SH2-SH3 adaptor proteins, ~30 Ras-GTPases and ~ 20 Rho-GTPases, and each set of GTPases is regulated by several dozens of GEFs and GAPs (HGNC; (Cherfils and Zeghouf, 2013). The diversity in these multi-layered and multi-family signaling proteins potentially supports a vast number of possible protein interactions and transduction cascades. A principle mechanism to achieve specificity and to customize the property of RTK signal transduction, elucidated mainly by studies in non-neuronal cells, is the assembly of signaling complexes where specific chains of protein interactions are organized by scaffolding proteins (Alberts, 2014). In the brain, however, whether RTK and Ras/Rho signaling are tailored to the needs and properties of different neuron types are unknown, in part due to a near absence of knowledge on their cellular expression patterns.

We found that members of the RTK family manifest significant differential expression among the 6 GABAergic populations (AUROC=0.74), suggesting that they might preferentially respond to different set of protein ligands. Downstream to RTKs, while multiple families of adaptor proteins (SH2 domain AUROC=0.69), GAPs (Rho-GAP AUROC=0.60; Arf-GAPs AUROC=0.65), kinases (AUROC: PI3Ks=0.61, MAPKKK=0.52, MAPKK=0.50, MAPK=0.57s) are more broadly expressed, the Ras and Rho signaling and regulatory proteins are differentially expressed among GTPs. Within the Ras family, 21 of the 32 members showed major enrichment in specific GTPs (AUROC=0.84). Different Ras family members might be activated by different upstream signals, have different cellular functions, and engage different downstream effectors (Buday and Downward, 2008; Mitin et al., 2005). This result suggests that GTPs might use Ras isoforms to relay distinct external inputs and trigger appropriate transcription programs and other effectors that mediate long term cellular changes.

Furthermore, both the Rho-GTPases and Rho-GEFs are differentially expressed. 37 of the 57 Rho-GEFs (AURPC=0.82) and 14 of the 19 Rho-GTPases (AUROC=0.72) are enriched in specific GTPs (Figure 5D). As different Rho isoforms are often activated by designated GEFs (Cook et al., 2014), our results suggest that differential expression of Rho signaling and regulatory components might provide the mechanism and capacity to maintain the diversity of GABAergic neuron morphology, connectivity, and to support different forms of neurite and synaptic motility and plasticity.

## 4. Transcription factor profiles register the developmental history and contribute to the maintenance of GTP phenotypes

### GABAergic neurons retain a transcription resume that registers their developmental history

The embryonic subpallium contains a developmental plan embedded in progenitors along ganglionic eminence whereby transcription cascades orchestrate the specification and differentiation of major clades (i.e. MGE, CGE) of cortical GABAergic neurons (Kepecs and Fishell, 2014; Nord et al., 2015) (Figure 7A-B). In response to early morphogen gradients, subpallium progenitors acquire a transcription program involving Gsx1/2, Zeb2, Ascl1 and Dlx1/2, which confers GABAergic fate. Subsequently, Nkx2.1 defines the MGE lineage and Coup-TFII the CGE lineage (Kepecs and Fishell, 2014; Nord et al., 2015). Along the MGE lineage, Lhx6 is a direct target of Nkx2.1 and acts in postmitotic neuronal precursors to trigger subsequent transcription programs that guide migration, differentiation and maturation (Du et al., 2008; Zhao et al., 2008). Downstream TFs include: Mafb, a early marker of MGE postmitotic percursors (McKinsey et al., 2013); Sox6, which regulates the positioning and maturation of PV and SST cells (Azim et al., 2009; Batista-Brito et al., 2008); and Satb1, an activity-modulated chromatin regulator required for the terminal differentiation and connectivity of interneurons (Close et al., 2012; Denaxa et al., 2012).

Along the CGE lineage, Coup-TFII, Sp8 and Prox1, Npas1, Npas3 contribute to the specification and differentiation of this major interneuron clade (Lodato et al., 2011; Ma et al., 2012; Miyoshi et al., 2015; Stanco et al., 2014). Several studies reported that these “embryonic TFs” are also expressed in subsets of mature cortical GABAergic neurons (Batista-Brito et al., 2008; Close et al., 2012; Miyazaki et al., 2012; Touzot et al., 2016). However, whether the expression patterns of these developmental TFs in mature cortex maintain their developmental history and the extent to which other TFs show similar “developmental continuity” are unclear.

By hierarchical and pair-wise comparison, we defined multiple sets of TFs that distinguish PV vs SST population (Figure 7F), and ChC vs PV, SST/nNOS vs SST/CR, and VIP/CR vs VIP/CCK cells (Figure 7G). Again, in cases where the developmental expression are reported there is a consistent developmental continuity of TF expression from embryonic precursors to mature neurons (e.g. ssbp2, Npas3, Bcl11) (Batista-Brito et al., 2008; Nikouei et al., 2016). In addition, by screening the Allen Atlas, we found that Klf5, Lmo4, Bcl11a (in SST cells), Pou3f2, Hey1 (in SST/nNOS cells) and Sox15 (in SST/CR cells) are expressed in embryonic ganglionic eminence neuronal progenitors and/or precursors (Figure S7B).

